# Dendritic cell PD-L2 restrains intratumoral CD8^+^ T cell immunity

**DOI:** 10.64898/2026.09.10.750646

**Authors:** Jinwoo Nah, Amanda Sun, Anushka Yadav, Adina Scheinfeld, Tamjeed Azad, Samuel Rose, Burkhard Becher, James W. Smithy, Michael A. Postow, Charlotte E. Ariyan, Yuri Pritykin, Chrysothemis C. Brown

**Affiliations:** Howard Hughes Medical Institute and Immuno-Oncology Program, Memorial Sloan Kettering Cancer Center, New York, NY, USA; Lewis-Sigler Institute for Integrative Genomics, Princeton University, Princeton, NJ, USA; Computational and Systems Biology Program, Memorial Sloan Kettering Cancer Center, New York, NY, USA; Institute of Experimental Immunology, University of Zurich, Zurich, Switzerland; Department of Medicine, Memorial Sloan Kettering Cancer Center, NY, USA; Weill Cornell Medical College, NY, USA; Department of Surgery, Memorial Sloan Kettering Cancer Center, NY, USA; Department of Computer Science, Princeton University, Princeton, NJ, USA; Immunology and Microbial Pathogenesis Program, Weill Cornell Medicine Graduate School of Medical Sciences, New York, NY, USA; Department of Pediatrics, Memorial Sloan Kettering Cancer Center, New York, NY, USA

## Abstract

Blockade of inhibitory PD-1 signaling on T cells is a cornerstone of cancer immunotherapy, with current strategies targeting PD-1 or its ligand PD-L1. However, PD-1 engages an alternative ligand, PD-L2, whose role in tumor immunity remains poorly defined. Here, we show that PD-L2 is upregulated on intratumoral CCR7⁺ conventional dendritic cells (cDCs) in both mouse and human melanoma. Using genetic mouse models enabling selective ablation of PD-L1 or PD-L2 in cDCs, we identify a division of labor between these ligands: PD-L1 controls the size of the progenitor CD8⁺ T cell pool in tumor-draining lymph nodes by modulating stem-like CD8⁺ T cells, whereas PD-L2 limits progenitor exhausted CD8^+^ T cell differentiation within the tumor microenvironment. Loss of PD-L2 in cDCs enhances cytotoxic CD8⁺ T cell responses and suppresses tumor growth, particularly in tumors enriched for CCR7⁺ cDC1s. Consistent with this, increased CCR7^+^ cDC abundance is associated with poor prognosis in human cancers. Spatial transcriptomic analyses reveal co-localization of CCR7⁺ cDC1s and Tpex within CCL19^hi^ niches, where cancer-associated fibroblasts serve as the predominant source of CCL19. Finally, intratumoral GM-CSF drives PD-L2 expression on CCR7⁺ cDCs, with Tpex and NK cells as major sources. Together, these findings establish cDC– associated PD-L1 and PD-L2 as spatially and functionally distinct checkpoints governing CD8⁺ T cell differentiation. Our results suggest that the abundance of CCR7⁺PD-L2⁺ cDC1s may guide the choice between anti–PD-1 and anti– PD-L1 therapies and support the development of PD-L2–directed blockade.

---

CD8^+^ T cells possessing cytotoxic “effector” function are essential for the elimination of infected and malignant cells^1,2^. Under conditions of persistent antigen encounter, such as in cancer or chronic infection, CD8^+^ T cells adopt a continuum of differentiation states, spanning from a reservoir of stem-like progenitor CD8^+^ T cells to transitory effector cells that ultimately progress toward terminally differentiated “exhausted” states with diminished cytotoxic capacity^3–8^.

The expression of ligand-dependent inhibitory receptors imposes critical checkpoints that restrain CD8^+^ T cell differentiation^9^. Among these, the PD-1 receptor and its ligand PD-L1 have emerged as key therapeutic targets in cancer immunotherapy^10,11^. PD-1 is expressed across the spectrum of activated CD8^+^ T cells, including stem-like progenitors as well as their more differentiated effector and exhausted counterparts^7,8^. Early models posited that PD-L1 expressed by tumor cells directly suppresses the activation and function of tumor-infiltrating PD-1^+^CD8^+^ T cells^12^. However, subsequent studies have revised this view, demonstrating that PD-1⁺ stem-like CD8⁺ T cells residing in tumor-draining lymph nodes (tdLNs) are the principal mediators of immune checkpoint blockade (ICB) responses^5,13,14^. In response to PD-L1 or PD-1 blockade, these quiescent cells undergo a proliferative burst, driving a new wave of tumor-infiltrating CD8^+^ T cells^5,15^. Within tdLNs, PD-L1^+^ type 1 conventional dendritic cells (cDC1s) regulate the maintenance and expansion of this progenitor pool, positioning cDC1s as critical intermediaries of checkpoint control^16–18^. At present, the mechanisms governing the transition of Tpex into functional effector populations within tumors are incompletely understood. Whether PD-1 signaling regulates the differentiation of progenitor exhausted CD8^+^ T (Tpex) cells following their entry into the tumor microenvironment remains unclear.

To date, investigation of PD-1 ligands has focused predominantly on PD-L1, and current immunotherapy primarily targets PD-1 or PD-L1^10,19^. However, PD-1 has an alternative ligand, PD-L2, which possesses substantially higher affinity^20–22^. In contrast to the well-defined immunoregulatory roles of PD-L1, the in vivo functions of PD-L2 remain comparatively underexplored^23,24^. Notably, PD-1 and PD-L1 blockade exhibit distinct efficacy and toxicity profiles, suggesting non-redundant roles for PD-L2^25,26^. Intriguingly, PD-L2 is also expressed by cDCs^20,27–29^, yet the role of cDC-derived PD-L2 in regulating CD8^+^ T cell-mediated immunity in lymph nodes and tumors is not fully known.

## RESULTS

### Tumor -associated CCR7^+^ cDCs upregulate PD-L2

To define the dynamic regulation of PD-L1 and PD-L2 expression on cDCs, we first profiled their expression across lymph node subsets at steady state (**Extended Data Fig. 1a**). Within the cDC1 compartment, PD-L1 expression was largely restricted to CCR7^+^ cells which represent activated or migratory cDCs^30,31^. In contrast, cDC2s exhibited broader PD-L1 expression across both CCR7^−^ and CCR7^+^ subsets (**Extended data Fig. 1b**). PD-L2 expression displayed a more restricted pattern, being confined almost exclusively to CCR7^+^ cDCs in both cDC1 and cDC2 lineages (**Extended data Fig. 1c**). Notably, whereas PD-L1 was universally expressed by CCR7⁺ cDC1s, the frequency of PD-L2⁺ cells within this population varied substantially across lymph nodes, ranging from ∼20% in axillary lymph nodes to ∼80% in gut-draining lymph nodes. These findings suggest that PD-L2 expression in cDC1s is dynamically regulated by tissue or lymph node-specific cues.

To determine how the tumor microenvironment (TME), both at baseline and following PD-1 blockade, shapes the dendritic cell (DC) landscape and the expression of inhibitory molecules, we performed single-cell RNA sequencing (scRNA-seq) on Lin⁻CD11c⁺MHCII⁺ DCs isolated from tumors, tumor-draining lymph nodes (tdLNs), and contralateral non-tumor draining lymph nodes (non-tdLNs) of mice bearing subcutaneous B16-OVA melanomas treated with anti–PD-1 or isotype control three days prior (**Fig. 1a**). Across non-tdLNs, tdLNs, and tumors, the overall composition of DC subsets was comparable between treatment groups (**Fig. 1b,c** and **Extended Data Fig. 2a-c**). Although canonical cDC1 and cDC2 subsets and their CCR7⁺ counterparts were transcriptionally similar between non-tdLNs and tdLNs, tumor-associated cDCs were distinct from their corresponding lymph node subset, suggesting imprinting by local cues within the TME (**Fig. 1d**). Notably, CCR7⁺ cDC populations were markedly enriched within tumors (**Fig. 1d**) – a surprising finding given that CCR7 upregulation is classically associated with egress of DCs from tissues in response to CCL19 gradients within lymph nodes^30,31^. To determine how the TME alters cDC states, we performed differential gene expression analysis comparing each tumor DC subset to its LN counterpart. Within both tumor CCR7^+^ cDC1 and cDC2s, *Pdcd1lg2* (encoding PD-L2) emerged among the most highly upregulated genes (**Fig. 1e**). In contrast, *Cd274* (PD-L1) expression was broadly detected across both tumor and lymph node CCR7⁺ cDCs, indicating a tumor-specific induction of PD-L2 (**Fig. 1f**). Flow cytometric analysis confirmed an increase in both the frequency of PD-L2⁺ cells and per-cell PD-L2 expression in tumor-associated CCR7⁺ cDC1s (**Fig. 1g**). Analysis across immune, stromal and tumor cells, further revealed that PD-L2 expression was largely restricted to cDC populations within the TME (**Extended Data Fig. 2d,e**). Importantly, scRNA-seq analysis on CD11c^+^MHCII^+^ dendritic cells and monocytes from human cutaneous melanoma demonstrated preferential expression of *PDCD1LG2* on CCR7⁺ cDCs (**Fig. 1h-j** and **Extended data Fig. 2f,g**), indicating that PD-L2 expression by tumor-associated CCR7^+^ cDCs is conserved across mouse and humans.

**Figure 1.**
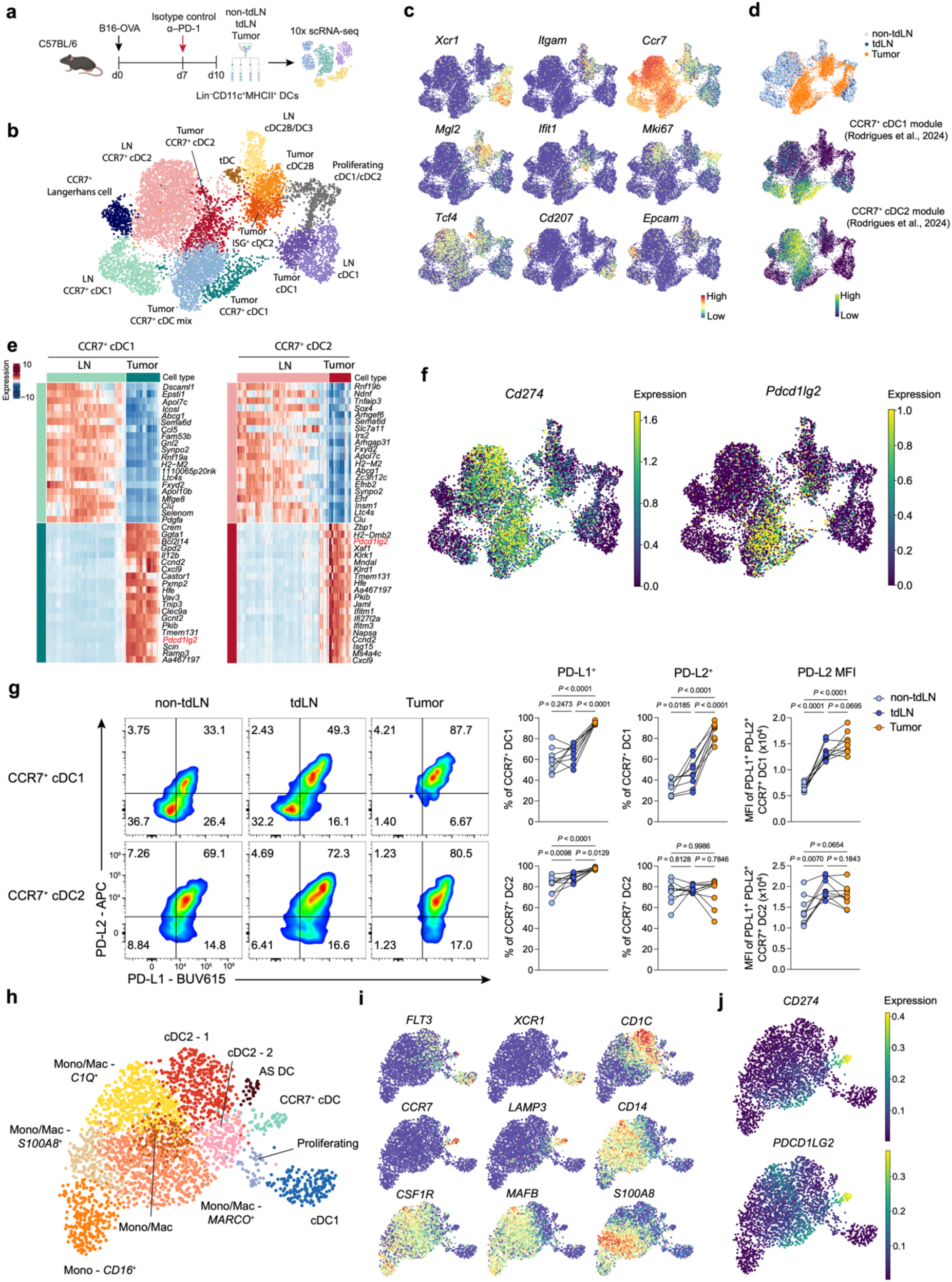
Tumor-associated CCR7⁺ cDCs upregulate PD-L2. **a***–***f**, scRNA-seq analysis of Lin⁻CD11c⁺MHCII⁺ DCs isolated from non-tumor-draining lymph nodes (non-tdLNs), tumor-draining lymph nodes (tdLNs) and tumors of B16-OVA-bearing mice treated with anti–PD-1 antibody or isotype control (n = 5 mice per group). **a**, Schematic of the experimental design. **b**, Uniform manifold approximation and projection (UMAP) of 9,660 cells colored by cluster annotation. **c**, UMAP colored by unimputed normalized expression of indicated marker genes. **d**, UMAP colored by tissue origin or CCR7⁺ cDC1 and CCR7⁺ cDC2 gene signature scores. **e**, Heatmaps showing scaled, imputed expression of the top 20 differentially expressed genes (one versus the rest, fold change (FC) > 1.5, adjusted *P* < 0.01) for tumor and lymph node CCR7⁺ cDC1 or cDC2 clusters. **f,** UMAP colored by unimputed normalized expression of PD-L1 (*Cd274*) and PD-L2 (*Pdcd1lg2*). **g**, Representative flow plots and summary graphs of PD-L1 and PD-L2 expression by CCR7⁺ cDC subsets from non-tdLNs, tdLNs and tumors of B16-OVA-bearing mice, analyzed 14 days post implantation (*n* = 9 mice). **h–j**, scRNA-seq on Lin(CD3, CD19, CD56)⁻CD11C⁺HLA-DR⁺ cells isolated from human melanoma. **h**, UMAP of 2,729 cells colored by cluster annotation. **i**, UMAP colored by unimputed normalized expression of the indicated marker genes. **j,** UMAP colored by imputed normalized expression of PD-L1 (*CD274*) and PD-L2 (*PDCD1LG2*). Panel **a** was created using BioRender; https://biorender.com. Data in **g** are pooled from two independent experiments and are shown as individual paired values, with lines connecting matched mice; one-way ANOVA (**g**). All *P* values are indicated on the corresponding graphs.

Collectively, these data identify CCR7⁺ cDCs as the dominant source of PD-L2 and establish that these cells accumulate within mouse and human melanoma, where local cues drive selective upregulation of PD-L2.

### cDC-derived PD-L1 restrains stem-like CD8⁺ T cell expansion and differentiation in tumor-draining lymph nodes

To dissect the overlapping and non-redundant roles of PD-1 ligands expressed by cDCs, we generated a conditional *Pdcd1lg2^fl^* allele to enable selective ablation of PD-L2 expression and bred either *Cd274^fl^* or *Pdcd1lg2^fl^* with *Clec9a^cre/cre^* mice to generate mice deficient in PD-L1 or PD-L2 expression by cDCs (hereafter *PD-L1^ΔDC^* or *PD-L2^ΔDC^* respectively). We observed complete deletion of the respective ligands in cDC1s and 50–70% of cDC2s (**Extended Data Fig. 3a,b**). Importantly, deletion of either ligand did not result in compensatory upregulation of the other (**Extended Data Fig. 3c**). To control for Clec9a deficiency in *Clec9a^cre/cre^* mice, we used *Clec9a^cre/cre^Cd274^fl/wt^* or *Clec9a^cre/cre^Pdcd1lg2^fl/wt^* littermate control mice in subsequent in vivo experiments. Compared with littermate wildtype mice (*Pdcd1lg2^wt/wt^*), mice with heterozygous loss of *Pdcd1lg2* showed comparable frequencies of PD-L2^+^ CCR7^+^ cDCs albeit with reduced PD-L2 expression levels on a per cell basis (**Extended Data Fig. 3d**). Under steady-state conditions, *PD-L1^ΔDC^* mice exhibited increased frequencies of PD-1⁺CD8⁺ T cells across various LNs and tissues relative to *Clec9a^cre/cre^Cd274^fl/wt^* control mice (**Extended Data Fig. 3e**), indicating a non-redundant role for PD-L1 in regulating CD8^+^ T cell homeostasis. In contrast, *PD-L2^ΔDC^* mice showed no detectable changes in the abundance of PD-1⁺CD8⁺ T cells across these sites, nor in CD4^+^ T cells, relative to *Clec9a^cre/cre^Pdcd1lg2^fl/wt^* control mice (**Extended Data Fig. 3f-h**). Although PD-L2 has been implicated in regulation of immune responses to the gut microbiota^28^, metagenomic sequencing of fecal samples revealed comparable bacterial communities between control and *PD-L2^ΔDC^* mice (**Extended Data Fig. 3i**), indicating that cDC-derived PD-L2 is dispensable for steady-state intestinal tolerance.

We next examined how cDC-derived PD-L1 and PD-L2 regulate anti-tumor immunity in B16-OVA tumors. To define the role of cDC PD-L1 across the continuum of tumor-specific CD8^+^ T cell differentiation – from stem-like progenitors in tdLN to effector and exhausted states within the tumor (**Fig. 2a**) – we adoptively transferred a low number of congenically marked naïve OT-I cells (5,000 cells) 6 hours before tumor implantation and analyzed their differentiation 10 days later (**Fig. 2b**). In *PD-L1^ΔDC^* mice, progenitor OT-I cells were markedly expanded within tdLNs, comprising Ly108^+^TIM3^−^CD62L^+^ cells, a subset previously associated with stem-like properties^32,33^ (hereafter referred to as stem-like T (T_SL_) cells), and Ly108^+^TIM3^−^CD62L^−^ progenitor exhausted T (Tpex) cells (**Fig. 2c** and **Extended Data Fig. 4a**). In addition, T_SL_ cells from *PD-L1^ΔDC^* mice exhibited increased expression of IL7R, a marker associated with memory potential^34^ (**Fig. 2d**). However, despite this expansion of the tdLN progenitor pool, the number and proportion of tumor-infiltrating Tpex or Teff/Tex OT-I cells was not altered in *PD-L1^ΔDC^* mice (**Extended Data Fig. 4b**). Similarly, we did not observe changes in the frequency of GZMB^+^, IL-2^+^, IFN-γ^+^ or TNF-α^+^ cells within these populations (**Extended Data Fig. 4c**). Consistent with the lack of enhanced intra-tumoral CD8^+^ T cell responses, ablation of PD-L1 on cDCs did not impact tumor growth (**Fig. 2e**).

**Figure 2.**
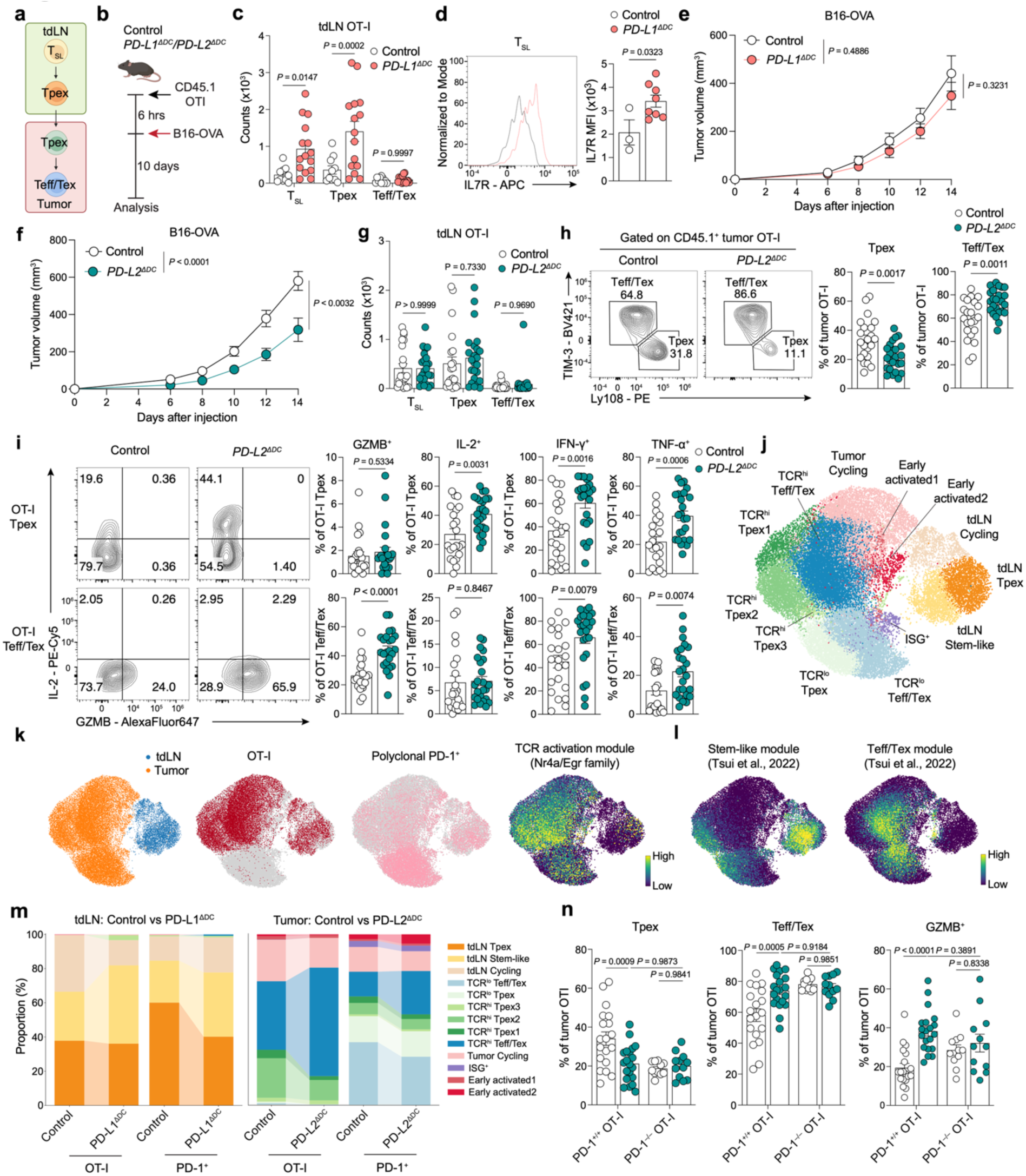
Dendritic cell PD-L1 and PD-L2 represent functionally and spatially distinct checkpoints in CD8 T cell differentiation. **a**, Schematic showing CD8 T cell differentiation across tdLNs and tumors. **b**, Schematic of the experimental design. **c**, Number of stem-like (T_SL_), progenitor exhausted (Tpex), and effector/exhausted (Teff/Tex) OT-I cells in tdLNs from *PD-L1^ΔDC^*(*n* = 15) and control (*n* = 10) mice. **d**, Representative flow plot and summary graph of IL7R expression on T_SL_ cells from *PD-L1^ΔDC^* (*n* = 8) and control (*n* = 3) mice. **e**, Tumor growth curves in *PD-L1^ΔDC^*(*n* = 9) and control (*n* = 8) mice following B16-OVA implantation without adoptive OT-I cell transfer. **f**, Tumor growth curves in *PD-L2^ΔDC^* (*n* = 16) and control (*n* = 15) mice following B16-OVA implantation without adoptive OT-I transfer. **g**. Number of T_SL_, Tpex, and Teff/Tex OT-I cells in tdLNs from *PD-L2^ΔDC^* (*n* = 24) and control (*n* = 22) mice. **h**, Representative flow plots and summary graphs of the frequency of Tpex (Ly108^+^TIM-3^−^) and Teff/Tex (Ly108^−^TIM-3^+^) cells among tumor-infiltrating OT-I cells from *PD-L2^ΔDC^*(*n* = 24) and control (*n* = 22) mice. **i**, Representative flow plots showing GZMB and IL-2 expression, and summary graphs of the frequency of GZMB^+^, IL-2^+^, IFN-γ^+^ and TNF-α^+^ cells among tumor-infiltrating OT-I Tpex and Teff/Tex cells from *PD-L2^ΔDC^* (n = 24) and control (n = 22) mice. **j–m**, scRNA-seq analysis of transferred OT-I cells and endogenous PD-1^+^CD8^+^ T cells isolated from tdLNs and tumors of B16-OVA-bearing *PD-L1^ΔDC^*, PD-L1 control, *PD-L2^ΔDC^*, and PD-L2 control mice. OT-I cells were pooled from 5, 6, 5 and 6 mice, respectively, whereas endogenous PD-1^+^CD8^+^ T cells were pooled from 4, 4, 6 and 5 mice, respectively. **j**, UMAP of 33,493 cells colored by cluster annotation. **k**, UMAP colored by tissue origin, cell-type origin, or TCR activation module score. **l**, UMAP colored by stem-like or Teff/Tex module scores, defined as in Tsui *et al*., 2022. **m**, Distribution of cell types as in (**j**) in tdLNs from *PD-L1^ΔDC^* and PD-L1 control mice (left) or tumors from *PD-L2^ΔDC^*and PD-L2 control mice (right). **n**, PD-1 deficient (PD-1^−/–^) or wildtype (PD-1^+/+^) OT-I cells were adoptively transferred into *PD-L2^ΔDC^*and control mice and analyzed as per the experimental scheme shown in (**b**). Summary graphs showing the frequency of Tpex, Teff/Tex, and GZMB^+^ cells among tumor-infiltrating OT-I cells from *PD-L2^ΔDC^* (n = 12 for PD-1^−/–^ OT-I, n = 20 for PD-1^+/+^ OT-I) and PD-L2 control (n = 11 for PD-1^−/–^ OT-I, n = 19 for PD-1^+/+^ OT-I) mice. Panel **a** and **b** were created using BioRender; https://biorender.com. Data in **c**, **e**, **f** are pooled from three independent experiments; Data in **d** are representative of two independent experiments; Data in **g–i** are pooled from six independent experiments; Data in **n** are pooled from seven independent experiments. Error bars: mean ± s.e.m.; two-way ANOVA (**c**, **e–g**, **n**), two-tailed unpaired t-test (**d–f**, **h**, **i**). All *P* values are indicated in the corresponding graphs.

Together, these findings indicate that cDC-mediated PD-L1/PD-1 signaling regulates the CD8⁺ T cell progenitor pool in tdLNs by restraining stem-like CD8⁺ T cell expansion and differentiation, but release of this checkpoint alone is insufficient to drive effector differentiation within tumors, implying the presence of additional inhibitory signals that limit this transition.

### cDC-derived PD-L2 restrains intratumoral Tpex-to-Teff/Tex differentiation

Given the selective upregulation of PD-L2 on tumor-associated CCR7⁺ cDC1s, we next asked whether PD-L2 regulates CD8⁺ T cell differentiation within tumors. In contrast to *PD-L1^ΔDC^* mice, *PD-L2^ΔDC^* mice showed significantly reduced B16-OVA tumor growth (**Fig. 2f**), indicating a non-redundant role for cDC-derived PD-L2 in anti-tumor immunity. Unlike *PD-L1^ΔDC^* mice, *PD-L2^ΔDC^* mice showed no expansion of T_SL_ and Tpex cells in the tdLNs following adoptive transfer of OT-I cells, as outlined above (**Fig. 2g**). Instead, PD-L2 deficiency promoted OT-I CD8⁺ T cell differentiation within tumors, with reduced Tpex frequency and a corresponding increase in Teff/Tex frequency (**Fig. 2h**). This shift was accompanied by increased frequencies of IL-2⁺ tumor-infiltrating OT-I Tpex cells and GZMB⁺, IFN-γ⁺ and TNF-α⁺ Teff/Tex cells (**Fig. 2i**), reflecting enhanced TCR activation and effector function^35^, consistent with the enhanced tumor control observed in *PD-L2^ΔDC^* mice.

To gain higher-resolution insight into regulation of CD8⁺ T cell differentiation by cDC-derived PD-L1 or PD-L2, we performed scRNA-seq on OT-I cells isolated from tdLNs and tumors of B16-OVA-bearing *PD-L1^ΔDC^* or *PD-L2^ΔDC^* mice, 10 days post transfer, as well as endogenous PD-1⁺CD8⁺ T cells from tumor-bearing mice without OT-I transfer (**Extended Data Fig. 5a**). This analysis revealed the full continuum of CD8⁺ T cell differentiation spanning tdLN stem-like and Tpex states to intratumoral Tpex and Teff/Tex populations (**Fig. 2j** and **Extended Data Fig. 5b-c**). Tumor OT-I cells and endogenous PD-1⁺CD8⁺ T cells preferentially distributed across distinct Tpex and Teff/Tex clusters that were distinguished by differential TCR signaling intensity, hereafter referred to as TCR^lo^ or TCR^hi^ Tpex and Teff/Tex (**Fig. 2k**). *Tcf7*⁺*Slamf6*⁺ progenitor cells within tdLNs spanned a cluster of *Sell*(CD62L)^+^*Il7r*^+^ cells enriched for stem-like module (tdLN Stem-like) and a cluster with lower stem-like module activity (tdLN Tpex). Within tumors, we identified both *Tcf7*⁺*Slamf6*⁺ Tpex states and Teff/Tex clusters enriched for cytotoxic and exhaustion-associated programs (**Fig. 2l** and **Extended Data Fig. 5d**).

Consistent with our earlier findings, the proportion of tdLN stem-like populations was increased in both OT-I and endogenous PD-1^+^CD8^+^ T cells from *PD-L1^ΔDC^* mice. By contrast, within tumors, *PD-L2^ΔDC^* mice showed a reduction in TCR^hi^ Tpex populations with a corresponding increase in TCR^hi^ Teff/Tex clusters in both OT-I and endogenous PD-1⁺CD8⁺ T cells (**Fig. 2m** and **Extended Data Fig. 5e**). Together, these data indicate that cDC-derived PD-L2, but not PD-L1, constrains the intratumoral Tpex-to-Teff/Tex transition, revealing a non-redundant role for PD-L2 in shaping CD8⁺ T cell differentiation within tumors.

To determine the therapeutic relevance of these findings, we performed checkpoint blockade using anti–PD-L1, anti–PD-L2, anti–PD-1 or isotype control antibodies (**Extended Data Fig. 5f**). Both PD-1 and PD-L2 blockade inhibited B16-OVA tumor growth, whereas PD-L1 blockade alone had no measurable effect (**Extended Data Fig. 5g**). Notably, PD-1 blockade produced a stronger tumor-control effect than PD-L2 blockade, consistent with the possibility that PD-1 integrates inhibitory signals from both PD-L1 and PD-L2^20^. These findings support a non-redundant role for PD-L2 in regulating anti-tumor immunity and raised the question of whether DC PD-L2–mediated suppression acts through PD-1.

PD-L2 has an alternative receptor, repulsive guidance molecule b (RGMb), which has been previously implicated in immune regulation^28,36^. To determine whether cDC-derived PD-L2 acts through RGMb or PD-1 to regulate tumor-specific CD8⁺ T cell differentiation, we first assessed RGMb expression following B16-OVA tumor implantation into *Rgmb*^GFP^ mice. RGMb (GFP) expression was not detected on CD8⁺ T cells in either tdLNs or tumors (**Extended Data Fig. 6a**), nor on transferred *Rgmb*^GFP^ OT-I cells in B16-OVA-bearing C57BL/6 mice (**Extended Data Fig. 6b**), suggesting that PD-1 may be the relevant CD8⁺ T cell receptor for PD-L2 in this model. Indeed, adoptive transfer of PD-1 deficient OT-I cells abrogated the phenotypic changes observed with wild-type OT-I cells in *PD-L2^ΔDC^* mice (**Fig. 2n**), confirming that cDC-derived PD-L2 restrains tumor-specific CD8⁺ T cell differentiation through PD-1 signaling.

Together, these results demonstrate that cDC-associated PD-L1 and PD-L2 function as spatially and temporally distinct checkpoints in CD8⁺ T cell differentiation: PD-L1 restrains expansion and differentiation of stem-like CD8^+^ T cells in tdLNs, whereas PD-L2 acts within tumors to limit Tpex differentiation into cytotoxic Teff/Tex cells.

### CCR7⁺ cDC1-rich TMEs define a PD-L2-regulated immunosuppressive niche

To determine whether cDC-derived PD-L2 regulates tumor control beyond B16-OVA, we evaluated additional tumor models in *PD-L2^ΔDC^* mice. cDC-specific PD-L2 deficiency did not alter tumor growth or OT-I CD8⁺ T cell responses in MC38-OVA or EO771-OVA tumors (**Fig. 3a,b** and **Extended Data Fig. 7a-f**). In contrast, *PD-L2^ΔDC^* mice implanted subcutaneously with ID8 ovarian cancer cells showed reduced tumor growth (**Fig. 3c**), recapitulating the phenotype observed with B16-OVA and consistent with previous work implicating PD-L2 in ovarian cancer immune evasion^37^. Given the striking enrichment of CCR7^+^ cDCs in B16-OVA tumors (**Fig. 1d**), we wondered whether the differential sensitivity of tumor models to cDC-specific PD-L2 deficiency reflected differences in the abundance of intratumoral CCR7⁺ cDCs. Consistent with this possibility, DC-PDL2 responsive B16 and ID8 tumors contained substantially higher frequencies of CCR7⁺ cDC1s relative to DC-PDL2 insensitive tumor models (**Fig. 3d**), suggesting that accumulation of CCR7⁺ cDC1s defines tumor contexts in which PD-L2-mediated regulation is most prominent.

**Figure 3.**
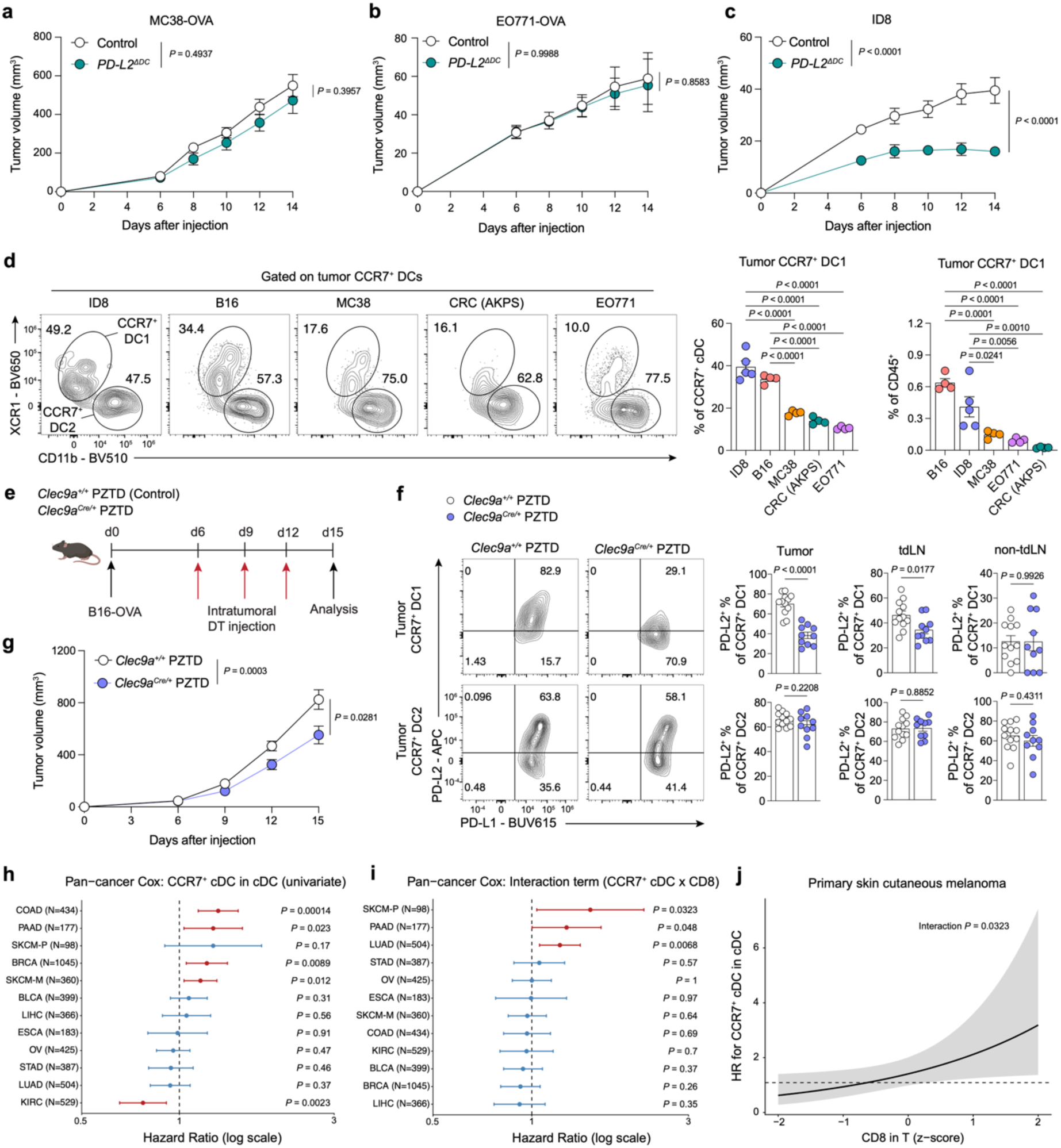
CCR7⁺ cDC1-rich TMEs define a PD-L2-regulated immunosuppressive niche. **a**, Tumor growth curves in *PD-L2^ΔDC^* (n = 6) and control (n = 9) mice implanted with MC38-OVA. **b**, Tumor growth curves in *PD-L2^ΔDC^*(n = 7) and control (n = 6) mice implanted with EO771-OVA. **c**, Tumor growth curve in *PD-L2^ΔDC^* (n = 9) and control (n = 8) mice implanted with ID8. **d,** Representative flow plots showing CD11b and XCR1 expression on tumor-associated CCR7⁺ cDCs across tumor models, and summary graphs showing the frequency of CCR7⁺ cDC1s among CCR7⁺ cDCs and total CD45⁺ immune cells (n = 4–5 mice per group). **e,** Schematic of intratumoral depletion of PD-L2⁺ cDCs using *Clec9a^cre/+^ PZTD* mice. **f,** Representative flow plots and summary graphs of PD-L1 and PD-L2 expression on CCR7⁺ cDC1 and cDC2 subsets in tumors, tdLNs and non-tdLNs from *Clec9a^cre/+^ PZTD* (n = 10) and *Clec9a*^+/+^ *PZTD* (n = 12) mice. **g,** Tumor growth curves in *Clec9a^cre/+^ PZTD* (n = 10) and *Clec9a*^+/+^ *PZTD* (n = 12) mice implanted with B16-OVA followed by DT treatment. **h–j**, Immune cell frequencies, including cDC subsets and CD8 T cells, were inferred across 12 TCGA cancer types using BayesPrism with a human melanoma scRNA-seq reference (Yang *et al*., 2025). **h,** Forest plot showing hazard ratios (HRs) for the frequency of CCR7⁺ cDCs among total cDCs across 12 TCGA cancer types by Cox regression survival analysis. **i**, Forest plot showing HRs for the CCR7⁺ cDC × CD8⁺ T cell interaction term from Cox regression across TCGA cancer types. **j,** HR estimate plot showing the association between CCR7⁺ cDC frequency and survival as a function of CD8⁺ T cell abundance z-score in primary skin cutaneous melanoma. Panel **e** was created using BioRender; https://biorender.com. Data in **a–c** are pooled from two independent experiments; Data in **d** are pooled from five independent experiments; Data in **f**, **g** are pooled from three independent experiments. Error bars: mean ± s.e.m.; two-tailed unpaired *t*-test (**a–c, f, g**), one-way ANOVA (**d**), two-way ANOVA (**a**-**c**, **g**). All *P* values are indicated in the corresponding graphs.

To directly test whether tumor-associated PD-L2⁺CCR7⁺ cDC1s are the principal PD-L2⁺ DC subset mediating suppression of CD8 anti-tumor immunity, we generated *Clec9a^cre/+^PZTD* (PD-L2–ZsGreen–TdTomato– Diphtheria toxin receptor) mice, in which PD-L2⁺CCR7⁺ cDC1s can be preferentially depleted by diphtheria toxin (DT) administration owing to reduced efficiency of cDC2 targeting by the heterozygous *Clec9a^cre^* allele. In B16-OVA–bearing *Clec9a^cre^*^/+^*PZTD* mice, low dose intratumoral DT administration selectively depleted PD-L2⁺CCR7⁺ cDC1s (**Fig. 3e,f**) and reduced tumor growth (**Fig. 3g**), indicating that intratumoral PD-L2⁺CCR7⁺ cDC1s restrain anti-tumor immunity.

Overall, these findings demonstrate that tumor-associated CCR7⁺ cDC1s restrain anti-tumor immunity through PD-L2–PD-1 signaling and suggest that variation in CCR7⁺ cDC1 abundance may shape the immunosuppressive landscape of the TME. Given that PD-L2 expression by tumor-associated CCR7⁺ cDCs was conserved across mouse and human melanoma, we next asked whether CCR7⁺ cDC abundance in human cancers was associated with adverse clinical outcomes. To more accurately infer CCR7⁺ cDC infiltration from TCGA bulk RNA-seq datasets, we applied BayesPrism^38^ to deconvolute cellular composition using a human melanoma-derived scRNA-seq reference^39^. Analysis of 12 TCGA tumor types^40^ revealed that an increased CCR7⁺ cDC frequency among cDCs was associated with worse prognosis in several cancers including colon adenocarcinoma (COAD), skin cutaneous melanoma (SKCM), pancreatic adenocarcinoma (PAAD) and breast invasive carcinoma (BRCA) (**Fig. 3h**). Cox regression analysis further demonstrated that the adverse association between CCR7⁺ cDC abundance and survival was dependent on CD8⁺ T cell abundance, most prominently in primary SKCM (SKCM-P), followed by PAAD (**Fig. 3i**). In these tumors, hazard ratio estimates showed that the negative prognostic effect of CCR7⁺ cDC enrichment increased progressively with higher CD8⁺ T cell abundance, whereas little or no effect was observed in tumors with low CD8⁺ T cell abundance (**Fig. 3j**). Further subdivision of human CCR7⁺ cDCs into CCR7⁺ cDC1 and CCR7⁺ cDC2 subsets was not possible as orthologous transcriptional signatures for these subsets have not yet been established.

Together, these findings establish that accumulation of immunosuppressive CCR7⁺ PD-L2^+^ cDC1s within the TME constrains CD8^+^ T cell-mediated tumor immunity.

### Spatially defined PD-L2⁺CCR7⁺ cDC1–Tpex niches within tumors

The enhanced intratumoral effector CD8^+^ T cell function in *PD-L2^ΔDC^* mice could reflect regulation of Tpex→Teff differentiation and/or reinvigoration of Tex cells by PD-L2 deficient CCR7^+^ cDC1s. To better understand the mechanism underlying PD-L2 regulation of intra-tumoral CD8^+^ T cell differentiation, we performed spatial transcriptomic profiling of non-tdLNs, tdLNs and tumors from B16-OVA-bearing mice that received OT-I cells (n = 6), together with tdLNs from tumor-bearing mice without OT-I transfer (n = 4), to establish CCR7^+^ cDC1 interactions within tumor and tdLN (**Extended Data Fig. 8a,b**). We used a fully custom 10x Xenium 472-gene panel to identify immune and stromal populations, with a particular focus on markers of DC subsets (**Supplementary Table 1**). We complemented this analysis by Visium HD profiling of selected adjacent tumor sections.

Cell segmentation and clustering of the 10x Xenium data captured the full spectrum of immune, stromal and tumor cell heterogeneity, with broadly comparable cell-type composition across individual samples (**Extended Data Fig. 8c-k**). Within tumor CD8^+^ T cells, we identified rare Tpex populations expressing canonical marker genes (*Cxcr5*, *Ccr6*, *Gpm6b*, *Penk*, and *Cd200*), and Teff/Tex populations expressing signature genes (*Havcr2*, *Pdcd1*, and *Prf1*) (**Extended Data Fig. 8j**). Among antigen-presenting cells (APCs), we identified CCR7^+^ cDC1s and CCR7^+^ cDC2s in both tdLNs and tumors, as well as other major DC subsets (**Fig. 4a** and **Extended Data Fig. 8g,k**). Consistent with our earlier observations, Xenium analysis showed that *Pdcd1lg2* expression was substantially higher in tumor-associated CCR7⁺ cDCs than in their LN counterparts (**Fig. 4b**). Notably, comparison across all tumor-infiltrating cell subsets showed that PD-L2 expression was highest in CCR7⁺ cDCs (**Fig. 4c**). We also identified high PD-L2 expression within Tpex cells, likely reflecting contamination from spatially overlapping PD-L2⁺CCR7⁺ cDCs (**Fig. 4c**).

**Figure 4.**
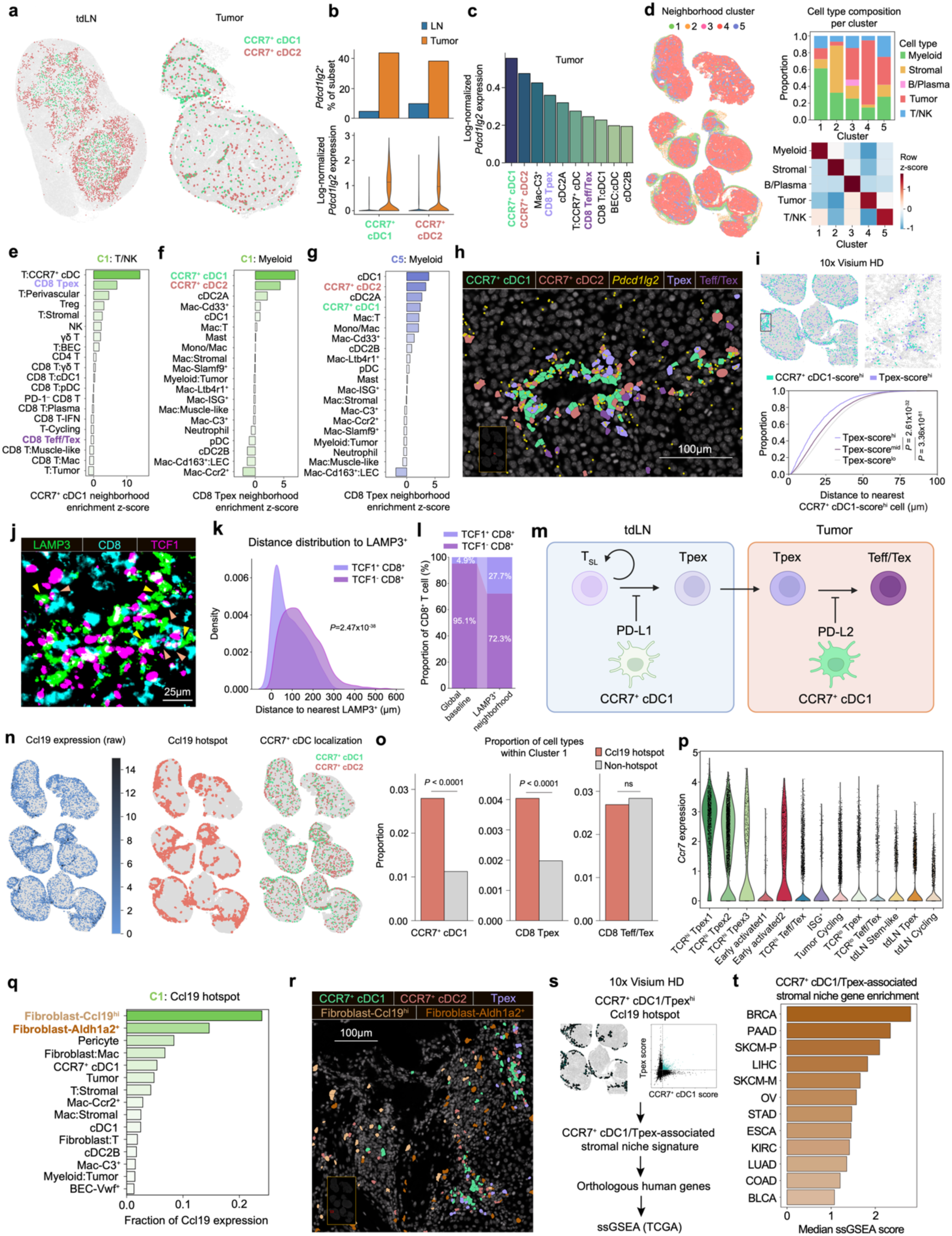
CCL19 marks spatially defined PD-L2⁺CCR7⁺ cDC1–Tpex niches within tumors. **a,** Representative spatial plots showing localization of CCR7^+^ cDC subsets in tdLN and tumor. **b**, Summary graphs of frequency of *Pdcd1lg2*^+^ and log-transformed library size-normalized *Pdcd1lg2* expression in CCR7^+^ cDC subsets from LNs and tumors. **c**, Summary graphs of log-transformed library size-normalized *Pdcd1lg2* expression in the highest-expressing cell types in tumors. **d**, Spatial plot, cell type composition and cell type enrichment heatmap of neighborhood clusters in tumors. **e**, Summary graph of enrichment z-scores of T/NK cell subsets around CCR7^+^ cDC1 neighborhoods within cluster 1. **f**, Summary graph of enrichment z-scores of myeloid subsets around CD8 Tpex neighborhoods within cluster 1. **g**, Summary graph of enrichment z-scores of myeloid subsets around CD8 Tpex neighborhoods within cluster 5. **h**, A representative spatial plot of a tumor region with CCR7⁺ cDC1, CCR7⁺ cDC2, Tpex and Teff/Tex cell localization, along with *Pdcd1lg2* expression. **i**, Visium HD spatial maps showing CCR7^+^ cDC1 signature-high and Tpex signature-high segmented cells, with a magnified view of the indicated region (top), and CDF plot showing the distance of Tpex signature-stratified segmented cells to the nearest CCR7^+^ cDC1 signature-high cell (bottom). **j**, Representative immunofluorescence imaging of human melanoma showing LAMP3⁺ DCs and TCF1⁻ and TCF1⁺ CD8⁺ T cells. **k**, A KDE plot showing the distance distribution of TCF1⁻ and TCF1⁺ CD8⁺ T cells to LAMP3⁺ DCs. **l**, Relative abundance of TCF1⁻ and TCF1⁺ CD8⁺ T cells in the global region or within a 20-µm-radius neighborhood centered on LAMP3⁺ DCs. **m**, Schematic delineating functionally and spatially distinct roles of cDC-derived PD-L1 and PD-L2 in CD8⁺ T cell differentiation. **n**, Spatial representation of *Ccl19* expression, identified Ccl19 hotspot regions and CCR7⁺ cDC localization in tumors. **o**, Summary graph showing the frequency of CCR7⁺ cDC1s, Tpex cells and Teff/Tex cells among all local cells in Ccl19 hotspot and non-hotspot regions from tumors. **p**, *Ccr7* expression in CD8⁺ T cell subsets from scRNA-seq data shown in Fig. 2j. **q**, Summary graph of the fraction of *Ccl19* expression contributed by indicated cell types within Ccl19 hotspots from cluster 1. **r**, A representative spatial plot of a tumor region with CCR7⁺ cDC subsets, Tpex cells and Ccl19-expressing cancer-associated fibroblasts (CAFs). **s**, Schematic of the analytic workflow. **t**, Summary graph showing CCR7⁺ cDC1/Tpex-associated stromal niche gene enrichment scores across 12 TCGA cancer types. Panel **m** was created using BioRender; https://biorender.com. Data in **b** are pooled from 16 LNs and 6 tumors. Data in **c–g**,**o**,**q** are pooled from 6 tumors. Data in **k** and **l** are from a region of interest (ROI) containing 57,733 total detected cells. Kolmogorov–Smirnov test (**i**), Mann–Whitney U-test (**k**), hypergeometric test (**o**). All *P* values are indicated in the corresponding graphs.

To define tumor neighborhoods in an unbiased manner, we applied a sliding-window approach to quantify local cell-type composition and identified five clusters representing different neighborhood states (**Fig. 4d** and **Extended Data Fig. 9a**). Clusters 1 and 2 corresponded to myeloid-enriched and stromal-enriched regions and were predominantly distributed in peritumoral areas. In contrast, clusters 3, 4 and 5 – B/plasma cell-, tumor- and T/NK-enriched regions – were mainly localized within intratumoral areas, revealing distinct anatomical organization of these neighborhood states (**Extended Data Fig. 9a,b**). CCR7⁺ cDC1s and cDC2s were most prominently enriched in cluster 1 (**Extended Data Fig. 9b**). Given these anatomically distinct neighborhood states with distinct cell type composition, we performed separate neighborhood enrichment analyses within each region to identify cell types spatially associated with CCR7⁺ cDC1s (**Fig. 4e-g** and **Extended Data Fig. 9c-e**). In cluster 1, where CCR7⁺ cDC1s were most abundant, their neighborhoods were strongly enriched for Tpex cells, but not Teff/Tex cells, indicating a preferential spatial association with Tpex cells (**Fig. 4e**). These neighborhoods showed limited enrichment of other myeloid subsets beyond CCR7⁺ cDC1s and CCR7⁺ cDC2s (**Extended Data Fig. 9d**), but were strongly enriched for blood endothelial cells (BECs) and pericytes (**Extended Data Fig. 9e**), consistent with previously described perivascular niches associated with CCR7⁺ cDCs ^41,42^. Reciprocal analysis of Tpex-centered neighborhoods confirmed that CCR7⁺ cDC1s were the predominant APC population enriched around Tpex cells in cluster 1 (**Fig. 4f**). Because Tpex cells were more strongly enriched in cluster 5 (a T/NK-enriched region) than in cluster 1, we also examined Tpex neighborhoods in cluster 5 regions. Here, CCR7⁺ cDC subsets were similarly highly enriched around Tpex cells along with cDCs (**Fig. 4g**). Spatial visualization illustrated preferential proximity between PD-L2-expressing CCR7⁺ cDCs and Tpex cells (**Fig. 4h**). Consistent with our Xenium-based analyses, Visium HD profiling of adjacent tumor sections showed spatial proximity between segmented cells with high CCR7^+^ cDC1 and Tpex signature scores derived from our scRNA-seq datasets (**Fig. 4i**).

Immunofluorescence analysis of human melanoma samples supported the translational relevance of these findings, revealing close spatial association between LAMP3⁺ cDCs (corresponding to human CCR7⁺ cDCs^29,43^) and TCF1⁺CD8⁺ T cells, with TCF1, encoded by *TCF7*, serving as a marker of progenitor-like CD8^+^ T cells (**Fig. 4j–l**).

Overall, these findings support a model in which tumor-associated CCR7⁺ cDC1s are spatially positioned to engage Tpex cells and restrain their differentiation into Teff/Tex cells, rather than directly suppressing the activation or effector function of terminally differentiated Teff/Tex cells (**Fig. 4m**).

### CCL19⁺ cancer-associated fibroblasts mark intratumoral CCR7⁺ cDC–Tpex niches

Classically, uptake of apoptotic tumor cells promotes cDC activation and differentiation into CCR7⁺ cDCs, which subsequently migrate to tdLNs along gradients of CCL19, a ligand for CCR7^31,44^. However, the accumulation of CCR7⁺ cDCs within specific tumor types suggested that local cues promote their retention within the TME. Analysis of the 10x Xenium dataset revealed discrete Ccl19^hi^ regions, which we defined as Ccl19 “hotspots”, closely associated with CCR7⁺ cDC localization (**Fig. 4n**). Within neighborhood cluster 1, CCR7⁺ cDC1s comprised a significantly higher proportion of cells in Ccl19 hotspots than in non-hotspot areas (**Fig. 4o**). Tpex, but not Teff/Tex cells, showed a similar enrichment pattern, indicating that CCR7⁺ cDC1s and Tpex cells are co-enriched within Ccl19-rich tumor regions. Notably, chemokine receptor analysis of our earlier scRNA-seq dataset revealed high *Ccr7* expression in TCR^hi^ Tpex clusters (**Fig. 4p**), suggesting that both cell types are recruited to these niches through CCR7-dependent chemotaxis and providing a potential basis for their co-localization.

To identify the cellular sources of CCL19, we quantified *Ccl19* expression across all cell types within Ccl19 hotspots. Within cluster 1 hotspots, *Ccl19*^hi^ fibroblasts represented the dominant source of Ccl19, followed by *Aldh1a2⁺* fibroblasts (**Fig. 4q**). These findings suggest that spatially and transcriptionally distinct cancer-associated fibroblast (CAF) populations establish CCL19-rich niches that may support the local accumulation of CCR7⁺ cDCs and Tpex cells within tumors (**Fig. 4r**).

To define the CCL19^+^ CAF–CCR7⁺ cDC1–Tpex niche at a transcriptome-wide level beyond the targeted gene panel captured by Xenium, we turned to the Visium HD data. Within Ccl19 hotspots, we identified regions enriched for CCR7⁺ cDC1s and Tpex cells, using signatures derived from our earlier scRNA-seq datasets. Genes upregulated in these regions defined a CCR7⁺ cDC1/Tpex-associated stromal niche signature (**Supplementary Table 2**). We then used an orthologous human gene signature to perform single-sample gene set enrichment analysis (ssGSEA) across 12 TCGA cancer types, assessing the clinical relevance of the CAF– CCR7⁺ cDC1–Tpex axis in human tumors (**Fig. 4s**). Enrichment scores were among the highest in PAAD and primary SKCM (**Fig. 4t**), the same tumor types in which CCR7⁺ cDC accumulation showed the strongest CD8⁺ T cell-dependent adverse prognostic association.

Collectively, these results identify a conserved CAF-associated program that marks CCL19-rich intratumoral CCR7^+^ cDC–Tpex niches across mouse and human tumors.

### Intratumoral GM-CSF drives PD-L2 upregulation on CCR7⁺ cDCs

A notable feature of tumor-associated CCR7⁺ cDC1s was their elevated expression of PD-L2 relative to lymph node CCR7⁺ cDC1s. To identify the microenvironmental cues driving PD-L2 expression, we leveraged a recently described “immune dictionary” of lymph node single-cell transcriptional responses to individual cytokines^45^. This identified GM-CSF as the top candidate regulator of both PD-L1 and PD-L2 expression in CCR7⁺ cDC1 and cDC2 subsets (**Fig. 5a**). Consistent with this prediction, in vivo GM-CSF blockade in B16-OVA-bearing mice resulted in a marked reduction in PD-L2 expression and a modest reduction in PD-L1 on tumor-associated CCR7⁺ cDCs, with minimal effects on tdLN cDCs (**Fig. 5b** and **Extended Data Fig. 10a,b**), suggesting that GM-CSF acts locally within the TME to promote PD-L2 expression. To identify the cellular source of GM-CSF, we analyzed B16-OVA-bearing *Csf2^icre-EGFP^R26^lsl-tdTomato^* mice. GM-CSF (GFP^+^ tdTomato^+^) expression was largely restricted to CD45⁺ immune cells within tumors, with PD-1^hi^CD8^+^ T cells and OVA-specific H-2Kb/SIINFEKL^+^CD8^+^ T cells representing a major source of GM-CSF, along with NK cells (**Fig. 5c,d** and **Extended Data Fig. 10c-e**). Among SIINFEKL^+^ CD8^+^ T cells, Tpex cells expressed higher levels of GM-CSF than Teff/Tex cells (**Fig. 5e**). Consistent with these findings, analysis of the 10x Xenium data revealed the highest levels of *Csf2* expression in Tpex cells among all cell types and prominent expression of the GM-CSF receptor (*Csf2rb*) by CCR7⁺ cDC1s (**Extended Data Fig. 10f**). Together, these findings identify GM-CSF as a local cue within the TME that promotes PD-L2 expression on CCR7⁺ cDCs.

**Figure 5.**
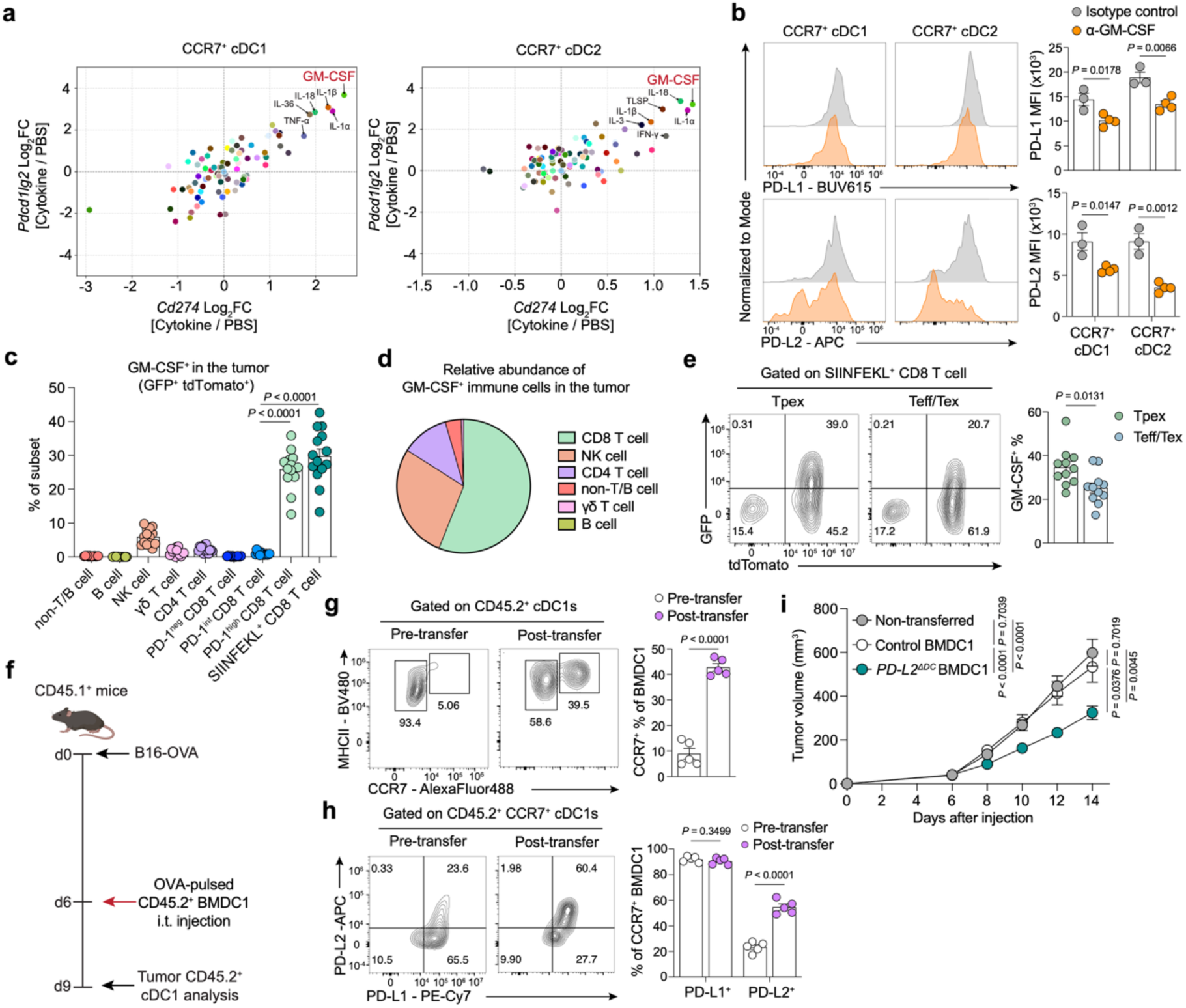
Intratumoral GM-CSF drives PD-L2 upregulation on CCR7⁺ cDCs. **a,** Fold change in *Cd274* and *Pdcd1lg2* expression in CCR7⁺ cDC subsets following stimulation with various cytokines, re-analyzed from the immune dictionary dataset (Cui *et al*., 2023). **b**, C57Bl/6 mice were implanted with B16-OVA, treated with anti–GM-CSF antibody or isotype control every 3 days, and analyzed on day 15. Representative flow plots and summary graphs of PD-L1 and PDL-2 expression by CCR7⁺ cDC subsets from anti–GM-CSF treated (n = 4) and isotype control treated (n = 3) mice. **c–e**, *Csf2^icre-EGFP^ R26*^lsl-tdTomato^ (FROG^Ai14^) mice were implanted with B16-OVA and analyzed on day 14. **c**, Summary graph of the frequency of GFP⁺tdTomato⁺ cells across the indicated immune cell populations (n = 14 mice). **d**, Pie chart showing the relative abundance of GM-CSF⁺ immune cell populations within tumors. **e**, Representative flow plots and summary graphs of the frequency of GM-CSF⁺ cells in SIINFEKL⁺ Tpex and Teff/Tex cells (n = 11 mice). **f,** Schematic of intratumoral transfer of OVA-pulsed CD45.2⁺ BMDC1s into B16-OVA-bearing CD45.1⁺ congenic mice. **g,** Representative flow plots and summary graph of CCR7 expression by XCR1^+^ BMDCs (BMDC1s) before and after transfer. **h,** Representative flow plots and summary graphs of PD-L1 and PD-L2 expression by CCR7⁺ BMDC1s before and after transfer. **i,** C57Bl/6 mice were implanted with B16-OVA tumors and left untreated or received intratumoral transfer of OVA-pulsed BMDC1s from control or *PD-L2^ΔDC^* mice on day 6. Tumor growth curves in mice with no BMDC1 transfer (n = 10), control BMDC1s (n = 9), and *PD-L2^ΔDC^* BMDC1s (n = 10). Panel **f** was created using BioRender; https://biorender.com. Data in **b**, **g**, **h** are representative of two independent experiments. Data in **c**, **d** are pooled from three independent experiments; data in **e, i** are pooled from two independent experiments. Error bars: mean ± s.e.m.; two-tailed unpaired t-test (**b, e, g**, **h**), one-way ANOVA (**c, i**), two-way ANOVA (**i**). All *P* values are indicated in the corresponding graphs.

The finding that GM-CSF drives PD-L2 expression on CCR7⁺ cDCs raised the possibility that a GM-CSF-rich tumor environment might limit the efficacy of DC-based immunotherapy. Indeed, intratumoral transfer of OVA-pulsed XCR1⁺CD103⁺ bone marrow-derived cDC1s (BMDC1s) into B16-OVA-bearing CD45.1⁺ congenic mice revealed upregulation of CCR7 expression on transferred BMDC1s and their retention within tumors (**Fig. 5f,g**). A substantial proportion of CCR7^+^ BMDC1s upregulated PD-L2 within the tumor, whereas PD-L1⁺ frequency remained largely unchanged, consistent with selective TME-driven induction of PD-L2 on CCR7⁺ cDCs (**Fig. 5h**). In line with their tumor-acquired immunosuppressive phenotype, control BMDC1s failed to promote tumor control, whereas transfer of PD-L2-deficient BMDC1s significantly inhibited tumor growth (**Fig. 5i**), indicating that TME-induced PD-L2 constrains the efficacy of DC-based immunotherapy.

Together these findings define a spatially organized circuit in which CCL19^+^ fibroblasts promote intra-tumoral CCR7^+^ cDC retention and their co-localization with CD8^+^ Tpex. In turn GM-CSF induces PD-L2 expression on CCR7^+^ cDCs, establishing a localized feedback loop that reinforces a PD-L2 checkpoint that constrains CD8^+^ T cell–mediated anti-tumor immunity (**Extended Data Fig. 10g**).

## DISCUSSION

Anti-PD1 blockade has been a cornerstone of cancer immunotherapy for over a decade^10,19^. Early models proposed that PD-L1 expressed by tumor cells directly suppresses the activation and effector function of tumor infiltrating PD-1^+^CD8^+^ T cells^12^. However, subsequent studies have revised this view, establishing PD-L1 on DCs as the critical regulator of CD8^+^ T cell-mediated tumor immunity and immunotherapy response, and demonstrating a role for CCR7^+^ cDC1s in regulating the reservoir of stem-like CD8^+^ T cells within tdLNs^13,16–18^. Here, we show that CCR7^+^ cDC1 regulation of CD8^+^ T cell immunity extends beyond the tdLN into the tumor, where the alternative PD-1 ligand PD-L2 restrains the differentiation of tumor-infiltrating Tpex into cytotoxic Teff/Tex states. Although some of our models and analyses capture both CCR7⁺ cDC1 and cDC2s, several lines of evidence including selective PD-L2 upregulation, enrichment in PD-L2-responsive tumors, PD-L2⁺ cDC1 depletion and spatial proximity to Tpex cells, implicate CCR7⁺ cDC1s as the principal mediators. Remarkably, despite high levels of PD-L1 expression by tumor-associated PD-L2^+^CCR7^+^ cDCs, PD-L1 cannot compensate for the loss of PD-L2, highlighting the non-redundant nature of PD-L1 and PD-L2 signaling on CD8^+^ T cells. Thus, cDC-associated PD-L1 and PD-L2 act as spatially and functionally distinct regulators of CD8⁺ T cell immunity, expanding the role of DCs beyond naïve T cell priming to active control of CD8⁺ T cell state transitions across lymph nodes and tumor tissues.

Recent studies have described CCR7⁺ cDCs–variably referred to as LAMP3^+^ DCs, mature DCs enriched in immunoregulatory molecules (mregDCs) or activated DCs–across multiple human cancers^29,39,43,46–50^. Spatial analyses place CCR7⁺ cDCs close to progenitor CD8⁺ T cells within intratumoral immune niches, and this CCR7⁺ cDC–Tpex spatial organization is conserved across multiple human tumor types, particularly within TLS-associated immune hubs^47,49,50^. Together with the association of CCR7⁺ cDC-rich niches with favorable responses to PD-1 blockade, these observations have supported the view that CCR7⁺ cDCs promote anti-tumor immunity. At first glance, this contrasts with our TCGA analysis, in which CCR7⁺ cDC signatures were associated with adverse overall survival in specific cancer types. However, because TCGA lacks information on immunotherapy exposure, this association may reflect baseline tumor biology rather than response to PD-1 blockade. Our findings suggest a context dependent role for CCR7^+^ cDCs: within Tpex hubs, CCR7^+^ cDC1s exert a predominantly immune-regulatory role, suppressing Tpex differentiation via PD-L2. Yet the PD-L2 dependent nature of this immunosuppression renders tumors harboring CCR7^+^ cDC-Tpex hubs particularly sensitive to anti-PD-1 or anti-PD-L2 blockade. Once PD-L2 inhibition is relieved, these same CCR7^+^ cDCs are poised to exert immune-stimulatory effects on CD8 Tpex within the niche.

We further identified CCL19⁺ CAF-rich tumor regions as stromal niches that bring CCR7⁺ cDC1s and Tpex cells into close proximity. A recent study established that CCL19-producing fibroblastic stromal cells organize CCR7^+^ cDC–CD8⁺ T cell niches in lymph nodes^51^. Spatial analysis of human non-small cell lung cancer (NSCLC) and head and neck squamous cell carcinoma (HNSCC) similarly identified CCL19^+^ CAFs near CCR7⁺ mature DCs, indicating that this stromal organization is conserved across species and tumor types^42,49,50^. Building on this, we show that GM-CSF is a key determinant of CCR7^+^ cDC PD-L2 expression, suggesting that intra-tumoral GM-CSF in part determines the immune-regulatory or immune-stimulatory character of CCR7^+^ cDC– Tpex niches.

In clinical practice, the choice between anti-PD-1 or anti-PD-L1 treatment has largely been determined by the first clinical trial to establish efficacy in specific tumor types, rather than direct comparative evidence which is largely lacking. However, in several tumor settings, anti–PD-1-based regimens have shown more consistent survival benefit than anti–PD-L1-based approaches, including gastric cancer, squamous non-small cell lung cancer (NSCLC), and triple negative breast cancer (TNBC)^25,52–55^. Although these comparisons are indirect and confounded by differences in trial design, they are compatible with the possibility that PD-L2–PD-1 interactions contribute to residual checkpoint suppression during PD-L1 blockade. Intriguingly, a recent analysis of PD-L2 expression in over 500 tumors suggested that high PD-L2 levels were associated with improved survival in patients that received immunotherapy, but not among immunotherapy-naïve patients^56^. These observations underscore the importance of rationally selecting checkpoint inhibitors based on the dominant regulatory pathways within a given tumor. While anti-PD-1 therapy targets both PD-L1 and PD-L2, it is also associated with increased immune-related adverse events, highlighting the need for more selective approaches^25,26^. Our findings suggest that the abundance of CCR7⁺ cDC1s, and by extension PD-L2 expression within the TME, may serve as a biomarker to guide the use of PD-L1-targeted or PD-L2-directed therapies, a prediction that warrants direct evaluation in treatment-stratified clinical cohorts. Conversely, in tumors lacking CCR7⁺ cDC infiltration, avoiding unnecessary blockade of PD-L2 may reduce toxicity without compromising efficacy.

Delineating the regulation and function of PD-L2 paves the way to new therapeutic approaches, including PD-L2 blockade or therapeutic targeting of the CCL19–GM-CSF–PD-L2 axis to enhance the efficacy of current immunotherapy or DC-based vaccination. Future studies will be required to determine how broadly this niche architecture is conserved across human cancers.

## ACKNOWLEDGEMENTS

We thank the Single-cell Analysis and Innovation Lab (SAIL) for single-cell genomic and spatial transcriptomic assays; the Molecular Cytology core (N. Fang, W. Kang, and E. Rosiek) for FFPE preparation and immunofluorescence imaging assays; C. Ariyan for B16 and B16-OVA tumor cell lines; S. Vardhana for MC38, MC38-OVA, EO771, and EO771-OVA tumor cell lines; K. Ganesh for AKPS tumor organoids; C. Reis e Sousa for the *Clec9a^cre/cre^* mice; S. Vardhana for the OT-I mice; J.F. Cloutier for the *Rgmb^GFP/+^* mice; J.P. Mizgerd for the *PZTD* mice; and the NIH Tetramer Core Facility (NIH Contract 75N93020D00005 and RRID:SCR_026557) for providing tetramers.

## AUTHOR CONTRIBUTIONS

J.N. and C.C.B. conceived the study, designed experiments, and wrote the manuscript. J.N. performed experiments, analyzed and interpreted data. Amanda S. designed and performed computational analyses for spatial transcriptomics data, with contributions from A.Y. and T.A.; A.Y. designed and performed computational analysis for scRNA-seq on T cells and human melanoma; Adina S. designed and performed computational analyses for scRNA-seq on B16-OVA DCs, with contributions from Amanda S. and A.Y.; S.R. designed and performed computational analysis for cytokine dictionary data; B.B. provided mice; M.P., J.S. and C.A. provided human tissues and intellectual expertise; Y.P. designed and supervised computational analyses of spatial transcriptomics data and scRNA-seq on B16-OVA DCs. C.C.B. supervised the research. All authors read and approved the manuscript.

## MATERIALS AND METHODS

### Mice

The conditional *Pdcd1lg2* allele, in which exon 2 is flanked by loxP sites, was generated by the Mouse Genetics Core Facility of Memorial Sloan Kettering Cancer Center (MSKCC) under protocol 90-12-033 approved by the MSKCC Institutional Animal Care and Use Committee. C57BL/6J mice (Jackson Laboratory, stock #000664) were used as zygote donors, and F1 females (C57BL/6J × CBA, stock #000656) served as recipients for embryo transfer. The 5′ loxP site was introduced by pronuclear microinjection of one-cell zygotes (collected from superovulated C57BL/6J females mated to C57BL/6J males) with a CRISPR–Cas9 cocktail comprising Cas9 protein (50–100 ng/µl; IDT), two crRNAs (25–50 ng/µl each; IDT) targeting *Pdcd1lg2* introns 1 and 2 to flank exon 2, tracrRNA (50–100 ng/µl; IDT) and a long single-stranded DNA (lssDNA) knock-in donor (1,663 nt; 5– 10 ng/µl; Megamer, IDT) in 1 mM Tris (pH 7.4). Injected zygotes were recovered in KSOM medium^57^ and transferred surgically into the oviducts of 0.5-d.p.c. pseudopregnant females. Founder (P) mice were genotyped by PCR for 5′ and 3′ loxP insertion. Multiple founders showed evidence of the 5’ loxP insertion. To introduce the second (3′) loxP site, zygotes were generated by in vitro fertilization using sperm from a 5′ loxP-positive founder male and wild-type C57BL/6J oocytes, electroporated as previously described^58^ with minor modifications. Briefly, 9–12 h after insemination, zygotes were washed in electroporation buffer (0.01% polyvinyl alcohol in Opti-MEM; Gibco) and electroporated in a platinum-electrode slide (Protech International) containing Cas9 protein (50–100 ng/µl; IDT), crRNA (25–50 ng/µl; IDT) and tracrRNA (50–100 ng/µl; IDT) with a single-stranded oligonucleotide (ssODN) donor (160 nt) using a Genome Editor electroporator (Protech International; three reciprocal pulse sets, six 3-ms pulses total, 97-ms intervals, 20 V). Electroporated zygotes were cultured in KSOM and transferred into pseudopregnant recipients. F1 Offspring were genotyped by PCR for both loxP sites. Mice positive for the inherited 5′ loxP and the newly introduced 3′ loxP were crossed to wild-type C57BL/6J mice; co-segregation of both loxP sites in F2 progeny confirmed the two sites were in cis, establishing the conditional *Pdcdl1lg2-*flox allele.

*Clec9a^cre/cre^*, *Rgmb^GFP/+^*, *Csf2^icre-EGFP^* (FROG; Fate-map and Reporter Of GM-CSF), *PZTD* (PD-L2–ZsGreen– TdTomato–Diphtheria toxin receptor), and OT-I (C57BL/6-Tg(TcraTcrb)1100Mjb/J) mice have been previously described^59–63^. C57BL/6J (CD45.2), *Cd274^fl/fl^* (B6.Cg-*Cd274^tm^*^3^*^.1Shr^*/J), PD-1^−/–^ (B6.Cg-*Pdcd1^tm^*^1^*^.1Shr^*/J), CD45.1 (B6.SJL-*Ptprc^a^ Pepc^b^*/BoyJ) and *R26^lsl-tdTomato^*(Ai14) mice were purchased from Jackson Laboratories. Generation and treatments of mice were performed under protocol 21-05-007, approved by the Sloan Kettering Institute (SKI) Institutional Animal Care and Use Committee. All mice were bred and maintained in the SKI animal facility under specific-pathogen-free (SPF) conditions in accordance with institutional guidelines and ethical regulations. Both male and female mice were included in the study. Age, litter-matched mice were used unless otherwise specified. All animals used in this study were experimentally naïve and had no prior history of experimental manipulation.

### Human specimens

Human tissue samples were obtained in accordance with national guidelines. Written informed consent was obtained from all patients, and the study was approved by the Institutional Review Board (IRB) at Memorial Sloan Kettering Cancer Center (IRB protocol #19-101). Patient information is provided in **Supplementary Table 3**. Tumor tissues were collected from surgical specimens after macroscopic examination by a pathologist. For each specimen, a tissue fragment was formalin-fixed and paraffin-embedded (FFPE) for histological analysis, and the remaining tissue was processed immediately to generate single-cell suspensions.

### Tumor cell lines and organoids

B16, B16-OVA, MC38, MC38-OVA, EO771, EO771-OVA, and ID8 cancer cell lines, as well as APC^−/–^Kras^LSL-G12D/+^p53^−/–^Smad4^−/–^ (AKPS) colon tumor organoids, were used in this study. B16 and B16-OVA were provided by C. Ariyan; MC38, MC38-OVA, EO771, EO771-OVA were provided by S. Vardhana; and AKPS colon tumor organoids were provided by K. Ganesh. ID8 cell line was purchased from Cytion. Unless otherwise specified, male mice were used for B16 and B16-OVA tumor experiments and female mice were used for implantation of the remaining cancer cells, consistent with the sex origin of these cell lines. Female mice were used for AKPS orthotopic implantation. Cancer cell lines were cultured and passaged in Dulbecco’s modified Eagle medium (DMEM) supplemented with 10% fetal bovine serum and 1% penicillin–streptomycin. AKPS tumor organoids were maintained in Advanced DMEM/F12 (Thermo Fisher Scientific) supplemented with 10mM HEPES (Corning), 2mM GlutaMAX (Thermo Fisher Scientific), 1mM N-Acetyle-L-cysteine (Sigma), B27 supplement with Vit. A (Thermo Fisher Scientific), 100 µg/ml Primocin (InvivoGen), 50 ng/ml mouse EGF (Invitrogen), 50 ng/ml mouse FGF-2 (Peprotech), 100 ng/ml mouse IGF-1(Peprotech), 500nM A83-01 (Tocris), 100 ng/ml mouse noggin (Peprotech).

### Tumor induction

Unless otherwise specified, 3 × 10⁵ cancer cells were suspended in 50 µl PBS and injected subcutaneously into the shaved right flank of mice. For EO771 breast cancer cell lines, 3 × 10⁵ cells were injected orthotopically into the fourth mammary fat pad of female mice. For orthotopic colon tumor experiments, 2 × 10⁵ AKPS tumor organoids were injected intracecally. Tumor volumes were measured using calipers and the volume was estimated using the formula: volume = (0.5 × length × width²).

### OT-I CD8 T cell isolation and adoptive transfer

Naïve OT-I cells were isolated from spleens of CD45.1 OT-I mice using a Naïve CD8α⁺ T Cell Isolation Kit (Miltenyi Biotec), according to the manufacturer’s instructions. For B16-OVA tumor experiments, 5 × 10³ naïve OT-I cells were suspended in 200 µl PBS and injected intravenously into recipient mice 6 h before tumor induction. For MC38-OVA and EO771-OVA tumor experiments, 1 × 10³ naïve OT-I cells were transferred. Transferred OT-I cells were analyzed in non-tdLNs, tdLNs and tumors 10 days after tumor induction.

### Antibody blockade experiments

For antibody blockade experiments, the following antibodies were used: anti-PD-1 (Bio X Cell, BE0146, clone RMP1-14), anti-PD-L1 (Bio X Cell, BE0101, clone 10F.9G2), anti-PD-L2 (Bio X Cell, BE0112, clone TY25), anti-GM-CSF (Bio X Cell, BE0259, clone MP1-22E9). Rat IgG2a (Bio X Cell, BE0089, clone 2A3) or rat IgG2b (Bio X Cell, BE0090, clone LTF-2) antibodies were used as isotype controls. Mice received 100 µg of blocking antibody or isotype control per injection intraperitoneally. For PD-1 pathway blockade, anti-PD-1, anti-PD-L1, anti-PD-L2 or isotype control antibodies were administered on days 6, 9 and 12 after B16-OVA tumor implantation. For GM-CSF blockade, anti-GM-CSF or isotype control antibodies were administered on days 0, 3, 6, 9 and 12 after B16-OVA tumor implantation.

### Tissue processing for flow cytometry and scRNA-seq

Mice were euthanized by CO₂ inhalation, and organs were harvested and processed as follows. Lymphoid organs, liver and lung were digested in RPMI 1640 supplemented with 5% fetal bovine serum (FBS), 1% L-glutamine, 1% penicillin–streptomycin, 10 mM HEPES, 1 mg/mg collagenase A (Sigma, 11088793001) and 1 U/ml DNase I (Sigma, 10104159001) for 45 min at 37°C with shaking at 250 rpm. For liver and lung samples, three 1/4-inch ceramic beads (MP Biomedicals, 116540034) were added per sample to aid tissue dissociation. Small and large intestines were removed, flushed with PBS and incubated for 15 min in PBS supplemented with 5% FBS, 1% L-glutamine, 1% penicillin–streptomycin, 10 mM HEPES, 1 mM dithiothreitol and 1 mM EDTA. Samples were then filtered through 100-µm strainers and centrifuged to isolate intraepithelial lymphocyte compartments. The remaining intestinal tissues were washed and digested for 30 min in digestion solution with ceramic beads. Digested samples were filtered through 100-µm strainers and centrifuged to remove collagenase-containing solution. Single-cell suspensions from tissue samples were washed once with 40% Percoll.

For tumor experiments, tumor-draining lymph nodes (tdLNs) and contralateral non-tdLNs were collected and processed as described above. For CD8 T cell phenotyping experiments, lymph nodes were mechanically dissociated, filtered through 100-µm strainers and centrifuged without enzymatic digestion. For isolation of tumor-infiltrating lymphocytes (TILs), tumors were minced into small fragments and digested for 45 min in digestion solution with ceramic beads. After digestion, samples were filtered through 100-µm strainers, washed and centrifuged. Cell pellets were resuspended in 4 ml of 40% Percoll and layered over 2 ml of 70% Percoll. Density-gradient centrifugation was performed at 800g for 20 min at 25°C with no acceleration or brake. Cells at the interphase were collected, washed, and centrifuged.

### Stimulation for intracellular cytokine staining

TILs were transferred to 96-well round-bottom plates, washed and centrifuged. Cells were resuspended in 200 µl cRPMI supplemented with 50 ng/ml PMA (Sigma), 1 µg/ml Ionomycin (Sigma), 5 µg/ml Brefeldin A (Biolegend) and 2 nM Monensin (Biolegend), and stimulated for 4 h at 37°C. After stimulation, cells were centrifuged and processed for surface and intracellular antibody staining.

### Flow cytometry

For flow cytometric analysis, dead cells were excluded by staining with LIVE/DEAD Fixable Zombie NIR in PBS for 10 minutes at 4°C with inclusion of anti-CD16/32 to block binding to Fc receptors, prior to cell-surface staining. Extracellular antigens were stained for 30 minutes at RT in staining buffer (2% FBS, 0.1% Na azide, in PBS, diluted 1:1 with Brilliant Violet (BD Biosciences) staining buffer. For experiments involving SIINFEKL– OVA tetramer staining, cells were washed and stained with tetramer for 30 min at RT in staining buffer before cell-surface staining. Where mCD1d tetramers were used, tetramers were added directly to the antibody cocktail and stained together with surface antibodies. For intracellular protein analysis, cells were fixed and permeabilized with Cytofix (BD Biosciences) and/or Ebioscience Foxp3 kit, per manufacturer instructions. Intracellular antigens were stained for 30min or overnight at 4°C in the respective 1x Perm/Wash buffer. The antibodies used for flow cytometry and FACS are listed in **Supplementary Table 4.** Unless otherwise stated, we used the following gatings: CCR7^+^ cDC1s: Lin(B220, CD90, TCRβ, NK1.1, Ly6C, CD88)^−^ CD11c^+^MHCII^+^EpCAM^−^CD207^−^XCR1^+^CD11b^−^CCR7^+^ , CCR7^+^ cDC2s: Lin^−^CD11c^+^MHCII^+^EpCAM^−^CD207^−^ XCR1^−^CD11b^+^CCR7^+^, tdLN OT-I T_SL_: TCRβ^+^CD8α^+^CD4^−^CD45.2^−^CD45.1^+^Ly108^+^TIM-3^−^CD62L^+^, tdLN OT-I Tpex: TCRβ^+^CD8α^+^CD4^−^CD45.2^−^CD45.1^+^Ly108^+^TIM-3^−^CD62L^−^, tumor OT-I Tpex: TCRβ^+^CD8α^+^CD4^−^CD45.2^−^ CD45.1^+^Ly108^+^TIM-3^−^, tumor OT-I Teff/Tex: TCRβ^+^CD8α^+^CD4^−^CD45.2^−^CD45.1^+^Ly108^−^TIM-3^+^. Example flow plots for CCR7^+^ cDC subsets and OT-I subsets are shown in **Extended Data Fig. 1** and **Extended Data Fig. 4**, respectively. Samples were acquired on a Cytek Aurora.

### scRNA-sequencing

#### Dendritic cell scRNA-sequencing from B16-OVA-bearing mice

B16-OVA cells (1 × 10⁶) were implanted subcutaneously into female C57BL/6 mice. Seven days after tumor implantation, mice were treated with 250 µg anti–PD-1 (n = 5) or isotype control antibody (n = 5). Three days later, tumors, tumor-draining lymph nodes and contralateral non-tumor-draining lymph nodes were pooled from 5 biological replicates and processed as described earlier. Cells were enriched for dendritic cells using the Dynabeads Mouse DC Enrichment Kit (Thermo Fisher) according to the manufacturer’s instructions. Cells were incubated with anti-CD16/32, and extracellular antigens were stained for 30 min at room temperature in sorting buffer (2% FBS, 2 mM EDTA, in PBS) and labelled with BioLegend TotalSeq-B Hashtag antibodies. Lin⁻(CD3⁻CD19⁻CD90⁻)CD64⁻Ly6C⁻CD11c⁺MHCII⁺ cells were then sort-purified using a FACSAria II cell sorter (BD Biosciences). Detailed gating strategies are provided in **Supplementary Fig. 1**. Cells were sorted into cRPMI, before being pelleted and resuspended in RPMI-2% FBS at a final concentration of 700–1,200 cells per µl. Cell viability was confirmed >80% with 0.2% (w/v) Trypan Blue staining (Countess II). Single cell gene expression analysis was performed with the 10x Genomics Chromium instrument following the user guide manual for 3′ v3 chemistry. In brief, cells were captured in droplets. Following reverse transcription and cell barcoding in droplets, emulsions were broken and cDNA purified using Dynabeads MyOne SILANE followed by PCR amplification per manual instruction. Approximately 30,000 cells were targeted for each sample. Samples were multiplexed together on one lane of 10x Chromium (using Hash Tag Oligonucleotides - HTO) following previously published protocol^64^. Final libraries were sequenced on Illumina NovaSeq S4 platform (R1 – 28 cycles, i7 – 8 cycles, R2 – 90 cycles). For sample demultiplexing, HTO sequencing data were aligned to the HTO barcodes, and UMIs were counted for each cell using CITE-seq-Count. Using a two-component K-means algorithm, we partitioned logged HTO counts into two distributions: background noise (lower mean) and positive tags (larger mean). Each droplet was then assigned to its source sample based on tags in the positive signal component. We classified droplets with multiple assignments as doublets and those with a single assignment as singlets. This analysis was performed using SHARP v0.1.1 (https://github.com/hisplan/sharp).

#### Dendritic cell scRNA-sequencing from human melanoma

Fresh tumor tissue was dissociated by manual mincing followed by an incubation of 30 min at 37°C in RPMI with liberase (0.83mg/ml; Sigma-Aldrich), alternated with 3 rounds of dissociation with a gentleMACS^TM^ dissociator (Miltenyi). After dissociation, cell suspensions were filtered with a 100 µm filter, and resuspended in ACK lysis buffer before washing with RPMI-5% FBS. Human DCs from melanoma were enriched with Dynabeads Human DC Enrichment kit (ThermoFisher) in accordance with manufacturer’s instructions. Cells were then washed with PBS and stained with LIVE/DEAD Fixable Ghost Dye Red 780 followed by cell surface antibodies. Lin(CD3, CD19, CD56)^−^CD11c^+^HLA-DR^+^ cells were then FACS-isolated on a FACS Aria II sorter (BD Biosciences) into RPMI-2% FBS for single cell RNA-seq. Detailed gating strategies are provided in **Supplementary Fig. 2**. The scRNA-seq libraries were prepared following the user guide manual (CG00052 Rev E) provided by the 10X Genomics and Chromium Single Cell 3′ Reagent Kit (v2). Briefly, samples were encapsulated in microfluidic droplets at a dilution of ∼70 cells/µl. Encapsulated cells were subjected to reverse transcription (RT) reaction at 53°C for 60 min. After RT step, the emulsion droplets were broken and barcoded-cDNA was purified with DynaBeads, followed by 14-cycles of PCR-amplification (98°C for 180 s; [98°C for 15 s, 67°C for 20 s, 72°C for 60 s] x 12-cycles; 72°C for 60 s). 50 ng of PCR-amplified barcoded-cDNA was fragmented with the reagents provided in the kit and purified with SPRI beads to obtain an average fragment size of 600 bp. Next, the DNA library was ligated to the sequencing adaptor followed by indexing PCR (98°C for 45 s; [98°C for 20 s, 54°C for 30 s, 72°C for 20 s] × 10 cycles; 72°C for 60 s). The resulting DNA library was double-size purified (0.6-0.8X) with SPRI beads and sequenced on an Illumina NovaSeq platform (R1 – 26 cycles, i7 – 8 cycles, R2 – 96 cycles). FASTQ files were processed using the Sequence Quality Control (SEQC) pipeline (Azizi et al., 2018) and reads were aligned to the human genome hg38.

#### CD8 T cell 10x Genomics Flex scRNA-sequencing from B16-OVA-bearing mice

For analysis of tumor-specific CD8⁺ T cell responses, CD45.1 OT-I cells were adoptively transferred into PD-L1 control (*Clec9a^cre/cre^*; *Cd274^fl/+^*), *PD-L1^ΔDC^* (*Clec9a^cre/cre^*; *Cd274^fl/fl^*), PD-L2 control (*Clec9a^cre/cre^*; *Pdcd1lg2^fl/+^*), and *PD-L2^ΔDC^* (*Clec9a^cre/cre^*; *Pdcd1lg2^fl/fl^*) mice. Six hours after transfer, mice were subcutaneously implanted with B16-OVA tumor cells. To analyze endogenous CD8 T cells, a separate cohort of mice was injected with B16-OVA cells without prior OT-I transfer. On day 10 post tumor induction, tdLNs and tumors of B16-OVA-bearing PD-L1 control (n = 6 for OT-I, n = 4 for PD-1^+^CD8^+^ T cells), *PD-L1^ΔDC^* (n = 5 for OT-I, n = 4 for PD-1^+^CD8^+^ T cells), PD-L2 control (n = 6 for OT-I, n = 5 for PD-1^+^CD8^+^ T cells), and *PD-L2^ΔDC^* mice (n = 5 for OT-I, n = 6 for PD-1^+^CD8^+^ T cells) were processed as described above. Cells from tdLNs and tumors were incubated with anti-CD16/32 and stained for extracellular markers in sorting buffer for 30 min at 4°C. Cells were labelled with BioLegend TotalSeq-C hashtag antibodies during this staining step for sample multiplexing. Cells were washed and resuspended in cRPMI containing SYTOX Blue (Invitrogen) to exclude dead cells. Live OT-I cells (TCRβ⁺CD8α⁺CD45.1⁺) or endogenous PD-1⁺CD8⁺ T cells (TCRβ⁺CD8α⁺CD44⁺PD-1⁺) were sorted directly into 500 µl fixation buffer consisting of 4% formaldehyde in 1× Fixation and Permeabilization Buffer (10x Genomics, PN-2000517) using a FACSAria II cell sorter (BD Biosciences). Detailed gating strategies are provided in **Supplementary Fig. 3**. Cells were fixed for 16–20 h at 4°C. Fixed cells were centrifuged at 850g for 5 min at room temperature, and fixation was quenched by resuspension in 500 µl 1× Quench Buffer (10x Genomics, PN-2000516). Samples were then supplemented with 0.1 volumes of Enhancer (10x Genomics, PN-2000482) and 10% glycerol and stored at −80°C. Samples were thawed at room temperature, centrifuged at 850g for 5 min and resuspended in 1 ml 0.5× PBS containing 0.02% BSA. Cell concentration and viability were assessed by AO–PI staining using a LUNA-FX7 Automated Cell Counter (Logos Biosystems). Cells were then processed per hybridization according to the 10x Genomics recommendations. Hybridization reactions were prepared using 80 µl hybridization mix and 20 µl Mouse WTA probes (10x Genomics, PN-2000510 or PN-2000718) and incubated at 42°C for 16–24 h. After hybridization, samples were diluted in Post-Hyb Wash Buffer and counted by AO–PI staining using a LUNA-FX7 Automated Cell Counter. For each experiment, equal numbers of cells from each hybridization were pooled to ensure equal sample contribution. Cell pools were then washed three times in Post-Hyb Wash Buffer for 10 min at 42°C. After washing, cells were resuspended in Post-Hyb Resuspension Buffer, filtered through a 30-µm filter (Miltenyi Biotec) and counted to determine the input required for Chromium X loading. GEM encapsulation was performed using the Chromium X system with Chip Q according to the 10x Genomics protocol and the manufacturer’s recommendations for targeted cell recovery. GEMs were recovered and processed according to the 10x Genomics workflow. After GEM processing, products were pre-amplified and indexed to generate sequencing libraries. Libraries were sequenced on an Illumina NovaSeq X using standard dual indexing and demultiplexing. Raw BCL files were processed using Cell Ranger.

### Single-cell RNA-seq computational analysis

#### Preprocessing of the 10x dendritic cell scRNA-seq

Data preprocessing was conducted using the scanpy package v1.9.3. Initial quality control steps and normalization were carried out separately for each sample. Cells were first required to have at least 100 detected genes. Within each sample, cells were further filtered on the basis of percentage of mitochondrial counts and total UMI counts, with sample-specific thresholds set by inspection of the count distributions. Potential doublets were identified using Scrublet (doublet score >= 0.2) and removed. For the tumor samples, an additional round of filtering removed a distinct low-count population of cells. Genes were filtered based on presence in at least 1.0% of cells within a given sample, and the union of genes passing this filter across all samples was retained for downstream analysis. Gene expression counts were then normalized with analytical Pearson residual normalization from scanpy, using a theta value of either 10 or 100 for each sample, according to its mean and variance distribution. After normalization, the samples were concatenated, and genes with non-finite (NaN) residual values were removed. This resulted in a dataset of 11,440 cells and 11,013 genes.

PCA was run with 100 components, a kNN graph was built using 30 neighbors, 90 PCs, and cosine metric, and Leiden clustering was performed at a resolution of 2.6 (selected by silhouette score), resulting in 35 clusters. Prior to differential expression analysis, the dataset was restricted to protein-coding genes (GENCODE vM23 annotation), yielding 10,510 genes. Cluster labels were then manually assigned based on the differential expression results, expression of canonical genes, and previously defined gene signatures^65^. Using these analyses, we identified 4 low QC clusters (low total counts or high mitochondrial counts) and 6 contaminant clusters (macrophage, Thetis cell, pDC, lymphoid, tumor and unknown minor cluster) which were removed from downstream analysis resulting in a preprocessed dataset. PCA and Leiden clustering were repeated on this dataset with the same parameters as above, resulting in 28 total clusters.

#### Preprocessing of the 10x human melanoma dendritic cell scRNA-seq

Barcodes were filtered based on the number of RNA-seq transcripts (>100 and <31,000), and the fraction of mitochondrial transcripts (<5%). Finally, any genes detected in <2 cells in the scRNA-seq data were discarded. After clustering (described in ‘Dimensionality reduction, cell clustering, and visualization’), analysis of QC metrics and differentially expressed genes, we identified 3 minor low QC clusters (low quality: clusters 12, 13, 14) which were excluded from downstream analyses. In total, 2,729 cells remained, with a median scRNA-seq library-size of 8,007 from 17,702 genes. Cluster labels were manually assigned and curated based on expression of canonical genes and differentially expressed genes identified by MAST analysis.

#### Preprocessing of the 10x CD8 T cell scRNA-seq

scRNA-seq and HTO FASTQ files were aligned to mm39 (Cell Ranger mouse reference genome GRCm39-2024-A) and HTO barcodes were counted and demultiplexed using Cell Ranger v9.0.1 multi to generate RNA and HTO count matrices for each sample. Each sample was further demultiplexed based on HTO counts using HTODemux function in Seurat v4.4.0. RNA count matrices for individual samples were first merged into a single count matrix. Next, barcodes classified as singlets from HTO demultiplexing were further filtered based on the fraction of mitochondrial transcripts (<10%) and any genes detected in <2 cells in the scRNA-seq data were discarded. After clustering (described in ‘Dimensionality reduction, cell clustering, and visualization’), analysis of QC metrics and differentially expressed genes, we identified 3 minor contaminant or low QC clusters (macrophage: cluster 21, B cell: cluster 22, low quality: cluster 20) which were excluded from downstream analyses. In total, 33,493 cells remained, with a median scRNA-seq library-size of 3,781 from 14,190 genes. Cluster labels were manually assigned and curated based on expression of canonical genes, differentially expressed genes identified by MAST analysis or expression of previously defined gene signatures for T cell states^32^.

#### Dimensionality reduction, cell clustering, and visualization

Following quality control filtering, raw counts were library-size normalized, log transformed (‘log-normalized’ expression values) and scaled using Scanpy v1.10.3^66^. A nearest-neighbor graph was constructed using the top 50 principal components (PCs) with 30 nearest neighbors. Clustering was performed using PhenoGraph^67^. After excluding contaminant or low QC clusters, the remaining clusters were re-plotted and visualized using UMAP^68^, computed from the nearest neighbor graph built by PhenoGraph.

#### Differential gene expression tests

Differentially expressed genes (DEGs) between groups of cells were identified with MAST^69^, performed using Seurat functions. MAST was run on the log-normalized expression values. In all tests, genes were only considered if they were detected in at least 1% of the cells in at least one of the two groups compared (min.pct=0.01, logfc.threshold=0). In one-vs-rest DE tests comparing multiple groups, each group was compared to all the cells from other groups. Specific DE comparisons are described in the results. DEGs were reported according to their log-fold change (>1.5) and adjusted p-value (<0.01). The top DEG markers were subsequently selected for each group, based on fold change and p-value.

#### Gene signature scores

The T cell signature scores (TCR activation, Stem-like, and Teff/Tex modules) were computed using the sc.tl.score_genes() function with default parameters from Scanpy. Nr4a1, Nr4a2, Nr4a3, Egr1, Egr2, and Egr3 were used as signature genes for TCR activation, based on their induction as immediate-early transcriptional responses downstream of TCR signaling^70–72^. Stem-like and Teff/Tex module scores were calculated using DEGs reported by Tsui et al. 2022^32^ for cluster 5 (CD62L^+^ T_PEX_) and clusters 0/1 (T_EX_1/2), respectively. CCR7^+^ cDC1 and cDC2 signature scores were calculated using subset-specific DEGs identified from the original scRNA-seq dataset^65^, defined by log-transformed fold change >1.5 and adjusted *P* < 0.01.

#### Data imputation for scRNA-seq data

MAGIC imputation^73^ was applied to the log-normalized expression values to de-noise and recover missing values. Imputed data were used only for visualization of gene expression on heatmaps or UMAPs, where specified.

### Intratumoral diphtheria toxin (DT) administration

*Clec9a^+/+^*; PZTD or *Clec9a^cre/+^*; PZTD mice received intratumoral injections of low-dose diphtheria toxin (DT; Sigma, D0564, 4ng per injection) to minimize effects in tumor-draining lymph nodes. DT was administered on days 6, 9, and 12 after B16-OVA tumor implantation.

### TCGA deconvolution and survival analysis

Bulk RNA-sequencing data and corresponding clinical annotations from The Cancer Genome Atlas (TCGA) were obtained using the TCGAbiolinks R package. For each cancer type, gene expression quantification files generated using the STAR–Counts workflow were downloaded and processed as raw count matrices. Gene symbols were assigned based on TCGA annotation metadata, and duplicated gene symbols were collapsed by summing counts. To maintain consistency across cohorts and match the preprocessing strategy used for the reference analysis, only genes detected in at least 10% of bulk tumor samples within each cancer type were retained for downstream deconvolution.

Cell-type deconvolution was performed using BayesPrism^38^ with a common single-cell RNA-sequencing reference^39^ across 12 TCGA cohorts; colon adenocarcinoma (COAD), pancreatic adenocarcinoma (PAAD), primary skin cutaneous melanoma (SKCM-P), metastatic skin cutaneous melanoma (SKCM-M), breast invasive carcinoma (BRCA), bladder urothelial carcinoma (BLCA), liver hepatocellular carcinoma (LIHC), esophageal carcinoma (ESCA), ovarian serous cystadenocarcinoma (OV), stomach adenocarcinoma (STAD), lung adenocarcinoma (LUAD), kidney renal clear cell carcinoma (KIRC). The reference consisted of annotated major tumor and immune cell compartments, including malignant cells, B cells, CD4 T cells, CD8 T cells, regulatory T cells, NK cells, monocytes/macrophages, cDC1s, cDC2s, CCR7^+^ cDCs (annotated as mature dendritic cells (mDCs) in the reference), endothelial cells and other non-malignant populations. To reduce potential bias from uneven cell numbers across reference populations, the single-cell reference was downsampled before deconvolution while preserving all annotated cell types. Malignant cells were designated as the tumor reference population during deconvolution, and inferred cell-type fractions were used for downstream analyses.

To compare relative CCR7^+^ cDC enrichment across cancer types, we calculated the fraction of CCR7^+^ cDCs within the cDC compartment:

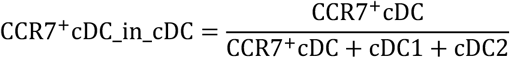

This metric was used to quantify relative skewing of the cDC compartment toward a CCR7^+^ cDC state, independent of overall cDC abundance. CD8 T cell enrichment within the T cell compartment was calculated as:

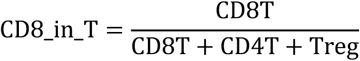

Overall survival time and event status were derived from TCGA clinical annotations using vital status, days to death and available follow-up information. Survival associations were assessed separately within each cancer type using Cox proportional hazards regression analysis. For each cohort, we evaluated the association of CCR7^+^ cDC enrichment and CD8 T cell enrichment with overall survival using univariable Cox models. Additionally, interaction models including CCR7^+^ cDC enrichment, CD8 T cell enrichment and their interaction term were used to test whether the prognostic effect of CCR7^+^ cDC enrichment varied according to CD8 T cell abundance. Hazard ratios with corresponding 95% confidence intervals were extracted from fitted models and summarized using forest plots for cross-cancer comparison of effect sizes.

### FFPE preparation

Tissues were dissected in ice-cold PBS and kept on ice before fixation. Samples were immersed in freshly prepared, ice-cold 10% formalin and fixed at 4°C. Specimens were mostly fixed for overnight, whereas larger tumor tissues were fixed for 2–4 days. Fixative volume was maintained at 10–20 times the tissue volume. After fixation, tissues were washed twice in PBS for 30 min each at 4°C and dehydrated in 70% ethanol for 30 min at room temperature or 4°C. Samples were stored in 70% ethanol at 4°C until paraffin processing. Tissues were dehydrated through graded ethanol washes, cleared in Sub-X clearing agent and infiltrated with paraffin. Samples were embedded using Leica HistoCore PEGASUS Parablocks and sectioned at 4 µm for downstream staining.

### Spatial transcriptomic 10x Xenium analysis

#### Tumor and lymph node profiling with 10x Xenium

Non-tdLNs, tdLNs and tumors were collected from B16-OVA-bearing mice that had received OT-I cell transfer (n = 6). In addition, tdLNs were collected from B16-OVA-bearing mice without OT-I transfer (n = 4). tdLN and tumor samples were pooled separately and processed for FFPE preparation and sectioned as described above. Xenium in situ gene expression profiling was performed on 4 µm formalin-fixed paraffin-embedded (FFPE) tissue sections using the Xenium platform (10x Genomics) according to the manufacturer’s protocol. Sections were mounted on Xenium slides, deparaffinized, and subjected to target retrieval and protease treatment to enable probe accessibility. A customized 472-gene panel, designed to identify immune, stromal and tumor cell populations, with a focus on DC subsets (**Supplementary Table 1**), was used for probe hybridization, followed by ligation and rolling circle amplification to generate localized amplicons. Cyclic fluorescence imaging was performed on the Xenium Analyzer to detect RNA targets at subcellular resolution. Cell segmentation staining was also performed with Xenium Multi-Tissue Stain. Primary image processing, transcript decoding, and cell segmentation were performed using Xenium Onboard Analysis (version 3.3.0.1; 10x Genomics) with default parameters. The analysis pipeline includes image registration across imaging cycles, spot detection, barcode decoding, and segmentation of individual cells based on nuclear and boundary signals, followed by assignment of detected transcripts to segmented cells. Output data consisted of spatially resolved gene expression profiles with associated cell-level annotations. CD45.1 SNP probes were included in the Xenium panel to detect transferred OT-I cells; however, these probes did not show specific signal across tissues, and OT-I cells could not be distinguished from endogenous CD8⁺ T cells in the Xenium data.

#### Data Preprocessing

Lymph node and tumor samples were processed separately. Within each tissue type, samples were concatenated and analyzed together. Cell transcripts were filtered to retain only those overlapping the nuclear segmentation for each cell. Following quality control filtering, raw counts were library-size normalized, log transformed (‘log-normalized’ expression values) and scaled using Scanpy v1.10.4. A nearest-neighbor graph was constructed using the top 30 principal components (PCs) with 15 nearest neighbors. Leiden clustering was performed, and clusters were manually assigned to broad cell type categories using expression of canonical genes. Clusters within each broad cell type category were merged, and clustering was performed again within each subset. Cluster labels were manually assigned and curated based on expression of canonical genes. Differential gene expression analysis was performed to identify genes enriched in each cluster using custom code. For this analysis, log fold change values of the mean expression of library-size normalized counts were calculated for each cluster vs all other cells. Multiple hypothesis test-corrected p-values were calculated using a Mann-Whitney U-test and the Benjamini-Hochberg procedure.

#### Sliding-window cell-type composition

A grid of circular sliding windows across the spatial data were defined as windows with a radius of 40 µm, a step size of 10 µm between windows, and at least 2 cells per window. Windows were added to cover any cells not included in the initial grid. K-means clustering was performed on the broad cell type composition of each window to identify five clusters. These clusters were labeled based on their enrichment of broad cell types. Cells were assigned the cluster label of the window with the closest center.

#### Neighborhood cell type enrichment

Cell-cell neighborhood connectivities were defined between cells with centers within 20 µm of each other. For each sample, cells and the corresponding connectivity graph were subset by neighborhood region. Within each sample-region subset, neighborhood enrichment by permutation test was performed as implemented in Squidpy v1.6.3^74^. Briefly, the observed number of connections between each pair of cell types was counted. Cell-type labels were then permuted 1000 times across cells while the connectivity graph was held fixed, and the number of connections between each cell type pair was recalculated for each permutation. For each cell type pair, a z-score was calculated by comparing the observed number of connections with the distribution obtained from the permutations. These z-scores were used as the neighborhood enrichment scores, with positive values indicating more connections than expected under the permutation null model and negative values indicating fewer connections than expected.

#### Ccl19 Hotspots

Ccl19-enriched cells were identified as cells with positive Ccl19 expression and smoothed Ccl19 expression within the top 30% of positive smoothed values. For each cell, smoothed Ccl19 expression was calculated as the mean Ccl19 expression among cells within a 100 µm radius. DBSCAN^75^ (eps= 100 µm, min_samples = 6) was applied to the spatial coordinates of Ccl19-enriched cells to identify spatially dense clusters. Cells not assigned to a cluster were classified as DBSCAN noise and excluded. A 20 µm-radius spatial expansion was added around each cell in the DBSCAN clusters to include cells nearby the Ccl19-enriched cells and overlapping buffer regions were merged. The Ccl19-enriched cells and nearby cells contained within the expanded regions comprised the Ccl19 hotspots.

### Spatial transcriptomic 10x Visium HD analysis

Tumor sections adjacent to those used for Xenium analysis were placed on the capture area of Visium Spatial Gene Expression for FFPE slides (10x Genomics). Slides were placed in a drying rack, incubated at 42°C for 3 h and then transferred to a desiccator overnight at room temperature. Slides were subsequently processed according to the 10x Genomics Visium FFPE workflow. Tissue sections were deparaffinized and stained with hematoxylin and eosin (H&E) to visualize tissue morphology. Library quality was assessed using the Agilent Bioanalyzer High Sensitivity DNA Kit (Agilent Technologies, 5067-4626). Libraries were sequenced on an Illumina NovaSeq X Plus platform using paired-end sequencing. Sequencing was performed with the following configuration: 28 bp for Read 1, containing the spatial barcode and UMI; 10 bp each for the i7 and i5 indexes; and 50 bp for Read 2, containing transcript sequences.

#### Data preprocessing

Samples were processed using Space Ranger v4.0.1, and the H&E-based cell-segmented output was used for downstream analyses. Following quality control filtering, raw counts were library-size normalized, log transformed, and scaled using Scanpy v1.10.4.

#### Gene signature scoring

T cell and dendritic cell signature scores were computed using the sc.tl.score_genes() function with default parameters in Scanpy. Gene signatures were derived from differential expression analyses performed in this study using CD8+ T cell scRNA-seq and dendritic cell scRNA-seq obtained from B16-OVA-bearing mice. Genes were included in each signature if they were upregulated with a log2 fold change >1.5 and an adjusted P value <0.05.

#### Stromal–cDC–Tpex niche analysis

Ccl19-enriched cells were identified in the same manner as the Xenium analysis described above, using the same parameters except that DBSCAN was performed with min_samples = 5. A 20 µm-radius spatial expansion was also applied as described above to define Ccl19 hotspots. Cells within Ccl19 hotspots with high values for both the Tpex and CCR7^+^ cDC1 signature scores, as defined by manual thresholds, were designated as high-scoring Ccl19 hotspot cells. Differential expression analysis was performed between high-scoring Ccl19 hotspot cells and cells outside any Ccl19 hotspot. Log fold changes were calculated from the mean expression in each group using library-size-normalized data. Statistical significance was assessed using a Mann-Whitney U-test, and P values were corrected for multiple hypothesis testing using the Benjamini-Hochberg procedure.

### Human melanoma Immunofluorescence (IF) analysis

Automated multiplex IF was conducted with the Leica Bond BX staining system. Paraffin-embedded tissues were sectioned at 4 µm and baked at 58°C for 1 hr. Slides were loaded in Leica Bond and IF staining was performed as follows. Samples were dewaxed at 72°C before being pretreated with EDTA-based epitope retrieval ER2 solution (Leica, AR9640) for 20 mins. at 100°C. The 5-plex antibody staining and detection was conducted sequentially. The primary antibodies against DC-LAMP/A488 (GT, 1:1000, R&D systems, AF4087), Xcr1/C594 (RT, 1:500,Abcam, ab317578), PDL-2/C430 (Rb, 1:500,Abcam, ab288298), CD8/A647 (Rb, 1/10, Roch Ventana, 790-4460) or TCF1/C543 (Rb, 1:500, 0.2ug/ml, Cell Signaling Technology, 2203) was incubated for 1h at RT followed by application of Leica Bond Polymer anti-rabbit HRP secondary antibody (included in Polymer Refine Detection Kit (Leica, DS9800)) for 8 min at RT. For the GT and Rat primary antibodies, a rabbit anti-Goat (Jackson ImmunoResearch 305-007-003) or a rabbit anti-Rat linker (Vector lab, BA4000) was incubated for 8 min before the application of the Leica Bond Polymer anti-rabbit HRP. After that, Alexa Fluor tyramide signal amplification reagents (Life Technologies, B40953, B40958) or CF® dye tyramide conjugates (Biotium, 92172, 96053, 92174) was used for detection. After each round of IF staining, Epitope retrieval was performed for denaturation of primary and secondary antibodies before another primary antibody was applied. After the run was finished, slides were washed in PBS and incubated in 5 µg/ml 4’,6-diamidino-2-phenylindole (DAPI) (Sigma Aldrich) in PBS for 5 min, rinsed in PBS, and mounted in Mowiol 4–88 (Calbiochem). Slides were kept overnight at -20°C before imaging. Slides were scanned on a Pannoramic Scanner (3DHistech, Budapest, Hungary) using a 20x/0.8NA objective.

Multiplex IF images were analyzed in QuPath-0.7.0 using StarDist-based cell segmentation. Nuclei were segmented from the DAPI channel using the ‘dsb2018_heavy_augment.pb’ StarDist model, with percentile normalization from 1–99%, a probability threshold of 0.5, and a detection pixel size of 0.5 µm. Segmented nuclei were expanded by 2 µm to approximate whole-cell boundaries, and shape and intensity measurements were extracted for each detection. Detections were generated within manually defined regions of interest. Cells were classified in QuPath based on marker-intensity thresholds into LAMP3⁺ cells, TCF1⁺CD8⁺ T cells, TCF1⁻CD8⁺ T cells. Cell centroid coordinates and classification labels were exported for downstream spatial analysis. Spatial analyses were performed in Python using custom scripts. Nearest-neighbor distances from each TCF1⁺CD8⁺ or TCF1⁻CD8⁺ T cell to the nearest LAMP3⁺ cell were calculated using Euclidean distance with SciPy cKDTree. Distributions were compared using two-sided Mann–Whitney U tests. For neighborhood-composition analysis, TCF1⁺CD8⁺ and TCF1⁻CD8⁺ T cells within 20 µm of LAMP3⁺ cells were aggregated to define pooled LAMP3-associated neighborhoods, and their composition was compared with the global CD8 T cell composition.

### Analysis of Immune Dictionary dataset

Raw count matrices for gene expression and hashtag detection for Cui *et. al.* (Immune Dictionary)^45^ were downloaded from GEO (GSE202186). The data was reprocessed to ensure accurate cell type and hashtag assignment, with a focus on myeloid lineage cells relevant for our study. To discriminate putative cell containing droplets from ambient RNA, we ran EmptyDrops^76^ with default parameters on each of 44 samples to estimate the likelihood of RNA within each droplet being derived from the ambient pool. Cell barcodes for droplets with a false discovery rate less than 0.01 were kept for analysis. Cell barcodes were then filtered again for only droplets that had a matching barcode detected in the corresponding hashtag sequenced sample denoting treatment information (generally > 99%). This produced a resulting matrix of 1,467,267 cells and 31,053 genes. Hashtags were assigned for each cell by taking the natural log + 1 of hashtag oligo counts in each cell divided by their geometric mean (centered log ratio) and then clustering the hashtags for each cell by kmeans with k = 2. For each cell if the hashtag with the highest expression was assigned to its own cluster, then that hashtag was assigned. Cells with 2 or more hashtags assigned or with no hashtag counts were called doublets or left unassigned, respectively. We additionally predicted putative doublet status by running DoubletDetection^77^ on raw count values for each sample individually (50 iterations, p_thresh=1e-7, voter_thresh=0.8). Cells that could not be assigned a hashtag were removed, as were cells with low counts (less than 350 UMIs or 250 genes) or low complexity (a value less than 0.8 log10(gene) / log10(UMI)). Values for the gene and UMI based cutoffs were derived from the position after the first mode in a histogram of all cells. After filtering the resulting gene expression matrix contained 595,769 cells.

Cytokine stimulations can induce dramatic changes in gene expression profiles that obscure cell type relationships during clustering. To ensure accurate cell type classification, a four step procedure was performed using both label transfer and manual annotation: 1) separate cells into lineage (stroma, myeloid, lymphoid) using a coarse human classification model; 2) cluster PBS treated cells; 3) use PBS treated cell type labels to train a classifier for projection to noisier cytokine treated cells; 4) cluster and manually annotate cells within sub-lineage based on marker gene expression. For cell type classification and model training, a logistic regression framework was used via the Celltypist package^78^.

To organize into distinct lineages, we used a publicly available cell type prediction model trained at a relatively coarse resolution^79^. Features for label transfer from a coarse human immune cell model (Immune_All_High from CellTypist model repo) were restricted to only one-to-one orthologous genes between human and mouse to support relevance across species (4,402 genes). To assign cell lineage, CellTypist was run with default parameters and classifications were assigned by majority voting of predicted labels within over-clustered data (knn = 30; 50 top PCs; leiden resolution 30). The assigned labels were then assigned to one of three broad lineages based on established hierarchies. This led to annotation of 107,971 myeloid cells, 484,447 lymphoid cells, and 2,499 non-hematopoietic cells (i.e. fibroblast, endothelial, epithelial).

We then focused on the myeloid lineage for sub-clustering. We trained a model for label transfer using the steady state PBS treated cells that could be projected to noisier cytokine treated data. For annotating the training data (PBS treated cells) we selected 3,000 highly variable genes using Scanpy (sc.pp.highly_variable_genes, flavor = ‘seurat_v3’, batch_key = processing batch) with only genes detected in at least 20 cells and excluding mitochondrial and ribosomal RNA genes and computed PCs with these features. Cells were clustered by Phenograph^67^ (k = 20, PCs = 50, Leiden partitioning algorithm with resolution 1) and assigned cell type labels based on marker genes. This included: pDC (*Tcf4*, *Bcl11a*), cDC1 (*Xcr1*, *Clec9a, Zbtb46*), migDC (*Ccr7*, *Flt3, Zbtb46*), langerhans cells (*Cd207*, *Epcam*), macrophage (*Maf*, *Spic*, *Rxra*, *Marco, Aif1*), neutrophil (*Cxcr2*, *Csf3r*), mast cell (*Kit*, *Cpa3*), and several clusters relating to monocyte derived cells or cDC2s (*Ccr2*, *Aif1, Csf1r, Irf4*) deemed macrophage/monocyte/DC (MMDC). We then selected for features informative to discriminating the cell types by computing marker genes between each coarse cluster using a Wilcoxon rank sum test against all other clusters. From this a maximum of 100 features (adjusted p < 0.01 & log fold-change > 0.5) were selected for each cell type, ranked by the test statistic, to use in the model (640 features). Cells within the PBS conditions were split into train/test at a 90/10 ratio to validate accurate predictions. Each cell type was down-sampled to 300 cells for training to balance classes. To ensure robustness to the down-sampling procedure, Celltypist models were trained with 10 different down-samplings and performance was assessed by classification accuracy on the held out test data. Across iterations, cell types were correctly assigned with > 97% median accuracy, and the model with the highest median accuracy was selected for use on cytokine treated cells. Cell type labels were then assigned to all cytokine treated cells by running CellTypist with the PBS cell model and using a majority vote approach with overclustering (knn = 30, PCs = 50, resolution = 25), using PCs computed with only features input to the CellTypist model.

Once cell type labels were assigned, cells were sub-clustered within the MMDC, Macrophage, and Langerhans/migDC groups to assign granular labels. Within each group, the top 2,000 HVGs were computed in Scanpy (flavor = ‘seurat_v3’) excluding mitochondrial genes and ribosomal RNA as well as any gene differentially expressed within the group with respect to cytokine treatment label (method = t-test; adjusted p value < .001; log_2_ fold-change > 1) to mitigate treatment influence during clustering. Each group was then clustered using Phenograph (50 PCs; knn = 20; resolution = 1) and cell state labels were assigned using canonical marker genes. The resulting processed and filtered expression matrix of myeloid cells contained 93,010 cells across 14 cell types. DC subsets were identified by expression of *Zbtb46* and distinguished from Thetis cells (TCs) by lack of *Rorc* expression. MigDCs were identified by *Ccr7* expression and comprised 3 clusters: migDC_apoptotic which had low counts and features and expressed high levels of mitochondrial genes indicating cell stress; CCR7^+^ cDC1 which expressed higher levels of *Laptm4b, Src,* and *Zdhhc14*; and CCR7^+^ cDC2 distinguished by higher levels of *Il2ra, Nrp2,* and *Capsl*. Expression values for *Cd274* and *Pdcd1lg2* were extracted from CCR7^+^ cDC1 and CCR7^+^ cDC2, and the mean expression of each gene was calculated for each cytokine group. Log2 fold changes relative to the PBS control condition were then calculated using a pseudocount of ε = 1 × 10⁻⁶ to avoid division by zero. The results were visualized as a scatter plot showing the log2 fold changes of both genes across all cytokine groups.

### Generation and adoptive transfer of BMDC1

Bone marrow cells were isolated from the indicated donor mice and cultured in six-well plates at 5 × 10⁵ cells/ml in 5 ml complete RPMI per well supplemented with recombinant mouse GM-CSF (2 ng/ml; PeproTech, 315-03) and recombinant human Flt3L-Fc (200 ng/ml; Bio X Cell, BE0098, clone Flt-3L-Ig (hum/hum)). Cells were incubated at 37°C with 5% CO₂ for 5 days. On day 5, 0.5 volumes of fresh culture medium (2.5 ml) were added to each well, and cells were cultured for an additional 4 days. On day 9, cultured cells were pulsed overnight with OVA (200 µg/ml; Sigma, A5503). The following day, CD11c⁺MHCII⁺CD103⁺XCR1⁺ cells were sorted using a FACSAria II cell sorter (BD Biosciences) and injected intratumorally into B16-OVA tumor-bearing recipient mice at 3 × 10⁵ cells per mouse.

### Statistics and reproducibility

Statistical analysis of data was performed using two-tailed unpaired *t*-tests, Kolmogorov–Smirnov tests, Mann–Whitney *U*-tests, Hypergeometric tests, one-way or two-way ANOVA adjusted for multiple-comparisons. Details of the number of replicates, sample size, significance tests and value and meaning of *n* for each experiment are included in the Methods or figure legends. Statistical analyses were performed using GraphPad Prism v.10 and v.11. *P* values of less than 0.05 were considered to indicate statistical significance, adjusted for multiple comparisons. All experiments were repeated at least twice as successful, independent experiments.

## DATA AND MATERIAL AVAILABILITY

Single cell and spatial datasets generated in this study are available through the Gene Expression Omnibus under accession number GSE(pending). This manuscript uses previously published scRNA-seq data from GSE262474, GSE188526, and GSE202186.

## CODE AVAILABILITY

Code for computational data analysis is available in GitHub at https://github.com/pritykinlab/DC-PD-L2-Tumor-Analysis.

## REFERENCES

1 Giles, J. R., Globig, A.-M., Kaech, S. M. & Wherry, E. J. CD8+ T cells in the cancer-immunity cycle. Immunity 56, 2231–2253 (2023).

2 Zhang, N. & Bevan, M. J. CD8+ T cells: foot soldiers of the immune system. Immunity 35, 161–168 (2011).

3 Wherry, E. J. T cell exhaustion. Nat. Immunol. 12, 492–499 (2011).

4 Blank, C. U. et al. Defining ‘T cell exhaustion’. Nat. Rev. Immunol. 19, 665–674 (2019).

5 Im, S. J. et al. Defining CD8+ T cells that provide the proliferative burst after PD-1 therapy. Nature 537, 417–421 (2016).

6 Utzschneider, D. T. et al. T cell factor 1-expressing memory-like CD8+ T cells sustain the immune response to chronic viral infections. Immunity 45, 415–427 (2016).

7 Miller, B. C. et al. Subsets of exhausted CD8+ T cells differentially mediate tumor control and respond to checkpoint blockade. Nat. Immunol. 20, 326–336 (2019).

8 Beltra, J.-C. et al. Developmental relationships of four exhausted CD8+ T cell subsets reveals underlying transcriptional and epigenetic landscape control mechanisms. Immunity 52, 825–841. e828 (2020).

9 Chen, L. & Flies, D. B. Molecular mechanisms of T cell co-stimulation and co-inhibition. Nat. Rev. Immunol. 13, 227–242 (2013).

10 Sharma, P. & Allison, J. P. The future of immune checkpoint therapy. Science 348, 56–61 (2015).

11 Sharpe, A. H. & Pauken, K. E. The diverse functions of the PD1 inhibitory pathway. Nat. Rev. Immunol. 18, 153–167 (2018).

12 Dong, H. et al. Tumor-associated B7-H1 promotes T-cell apoptosis: a potential mechanism of immune evasion. Nat. Med. 8, 793–800 (2002).

13 Connolly, K. A., et al. A reservoir of stem-like CD8+ T cells in the tumor-draining lymph node preserves the ongoing antitumor immune response. Sci. Immunol. 6, eabg7836 (2021).

14 Sade-Feldman, M. et al. Defining T cell states associated with response to checkpoint immunotherapy in melanoma. Cell 175, 998–1013. e1020 (2018).

15 Huang, Q. et al. The primordial differentiation of tumor-specific memory CD8+ T cells as bona fide responders to PD-1/PD-L1 blockade in draining lymph nodes. Cell 185, 4049–4066. e4025 (2022).

16 Schenkel, J. M. et al. Conventional type I dendritic cells maintain a reservoir of proliferative tumor-antigen specific TCF-1+ CD8+ T cells in tumor-draining lymph nodes. Immunity 54, 2338–2353. e2336 (2021).

17 Oh, S. A. et al. PD-L1 expression by dendritic cells is a key regulator of T-cell immunity in cancer. *Nat*. Cancer 1, 681–691 (2020).

18 Mayoux, M. et al. Dendritic cells dictate responses to PD-L1 blockade cancer immunotherapy. Sci. Transl. Med. 12, eaav7431 (2020).

19 Topalian, S. L., Drake, C. G. & Pardoll, D. M. Immune checkpoint blockade: a common denominator approach to cancer therapy. Cancer Cell 27, 450–461 (2015).

20 Latchman, Y. et al. PD-L2 is a second ligand for PD-1 and inhibits T cell activation. Nat. Immunol. 2, 261–268 (2001).

21 Youngnak, P. et al. Differential binding properties of B7-H1 and B7-DC to programmed death-1. Biochem. Biophys. Res. Commun. 307, 672–677 (2003).

22 Philips, E. A. et al. The structural features that distinguish PD-L2 from PD-L1 emerged in placental mammals. J. Bio. Chem. 295, 4372–4380 (2020).

23 Keir, M. E., Butte, M. J., Freeman, G. J. & Sharpe, A. H. PD-1 and its ligands in tolerance and immunity. Annu. Rev. Immunol. 26, 677–704 (2008).

24 Wang, Y. et al. Evolving landscape of PD-L2: bring new light to checkpoint immunotherapy. Br. J. Cancer 128, 1196–1207 (2023).

25 Wang, X. et al. Effectiveness and safety of PD-1/PD-L1 inhibitors in the treatment of solid tumors: a systematic review and meta-analysis. Oncotarget 8, 59901 (2017).

26 Sonpavde, G. P., Grivas, P., Lin, Y., Hennessy, D. & Hunt, J. D. Immune-related adverse events with PD-1 versus PD-L1 inhibitors: a meta-analysis of 8730 patients from clinical trials. Future Oncol. 17 (2021).

27 Tseng, S.-Y. et al. B7-DC, a new dendritic cell molecule with potent costimulatory properties for T cells. J. Exp. Med. 193, 839–846 (2001).

28 Park, J. S. et al. Targeting PD-L2–RGMb overcomes microbiome-related immunotherapy resistance. Nature 617, 377–385 (2023).

29 Maier, B. et al. A conserved dendritic-cell regulatory program limits antitumour immunity. Nature 580, 257–262 (2020).

30 Ohl, L. et al. CCR7 governs skin dendritic cell migration under inflammatory and steady-state conditions. Immunity 21, 279–288 (2004).

31 Förster, R., Davalos-Misslitz, A. C. & Rot, A. CCR7 and its ligands: balancing immunity and tolerance. Nat. Rev. Immunol. 8, 362–371 (2008).

32 Tsui, C. et al. MYB orchestrates T cell exhaustion and response to checkpoint inhibition. Nature 609, 354–360 (2022).

33 Gago da Graça, C., et al. Stem-like memory and precursors of exhausted T cells share a common progenitor defined by ID3 expression. Sci. Immunol. 10, eadn1945 (2025).

34 Kaech, S. M. et al. Selective expression of the interleukin 7 receptor identifies effector CD8 T cells that give rise to long-lived memory cells. Nat. Immunol. 4, 1191–1198 (2003).

35 Kaech, S. M. & Cui, W. Transcriptional control of effector and memory CD8+ T cell differentiation. Nat. Rev. Immunol. 12, 749–761 (2012).

36 Xiao, Y. et al. RGMb is a novel binding partner for PD-L2 and its engagement with PD-L2 promotes respiratory tolerance. J. Exp. Med. 211, 943–959 (2014).

37 Miao, Y. R. et al. Neutralization of PD-L2 is Essential for Overcoming Immune Checkpoint Blockade Resistance in Ovarian Cancer. Clin Cancer Res 27, 4435–4448 (2021).

38 Chu, T., Wang, Z., Pe’er, D. & Danko, C. G. Cell type and gene expression deconvolution with BayesPrism enables Bayesian integrative analysis across bulk and single-cell RNA sequencing in oncology. Nat. Cancer 3, 505–517 (2022).

39 Yang, J. et al. Mature and migratory dendritic cells promote immune infiltration and response to anti-PD-1 checkpoint blockade in metastatic melanoma. Nat. Commun. 16, 8151 (2025).

40 Weinstein, J. N. et al. The cancer genome atlas pan-cancer analysis project. Nat. Genet. 45, 1113–1120 (2013).

41 Di Pilato, M. et al. CXCR6 positions cytotoxic T cells to receive critical survival signals in the tumor microenvironment. Cell 184, 4512–4530. e4522 (2021).

42 Zitti, B. et al. Positioning and reversible suppression of CCR7+ dendritic cells in perivascular tumor niches shape cancer immunity. Immunity 59, 161–176. e112 (2026).

43 Zhang, Q. et al. Landscape and dynamics of single immune cells in hepatocellular carcinoma. Cell 179, 829–845. e820 (2019).

44 Roberts, E. W. et al. Critical role for CD103+/CD141+ dendritic cells bearing CCR7 for tumor antigen trafficking and priming of T cell immunity in melanoma. Cancer Cell 30, 324–336 (2016).

45 Cui, A. et al. Dictionary of immune responses to cytokines at single-cell resolution. Nature 625, 377–384 (2024).

46 Cheng, S. et al. A pan-cancer single-cell transcriptional atlas of tumor infiltrating myeloid cells. Cell 184, 792–809. e723 (2021).

47 Magen, A. et al. Intratumoral dendritic cell–CD4+ T helper cell niches enable CD8+ T cell differentiation following PD-1 blockade in hepatocellular carcinoma. Nat. Med. 29, 1389–1399 (2023).

48 Ma, T. et al. Pan-Cancer Analyses Refine the Single-Cell Portrait of Tumor-Infiltrating Dendritic Cells. Cancer Res. 85, 3596–3613 (2025).

49 Chen, J. et al. Human lung cancer harbors spatially organized stem-immunity hubs associated with response to immunotherapy. Nat. Immunol. 25, 644–658 (2024).

50 Mattiuz, R. et al. Dendritic cells control tertiary lymphoid structure development and maintenance in cancer. Science 393, eady1678 (2026).

51 Alouche, N. et al. Homeostatic mature dendritic cells instruct fibroblast specialization via Notch2 signaling to establish T cell niches. Immunity (2026).

52 Duan, J. et al. Use of Immunotherapy With Programmed Cell Death 1 vs Programmed Cell Death Ligand 1 Inhibitors in Patients With Cancer: A Systematic Review and Meta-analysis. JAMA Oncol. 6, 375–384 (2020).

53 Miles, D. et al. Primary results from IMpassion131, a double-blind, placebo-controlled, randomised phase III trial of first-line paclitaxel with or without atezolizumab for unresectable locally advanced/metastatic triple-negative breast cancer. Ann. Oncol. 32, 994–1004 (2021).

54 Cortes, J. et al. Pembrolizumab plus chemotherapy versus placebo plus chemotherapy for previously untreated locally recurrent inoperable or metastatic triple-negative breast cancer (KEYNOTE-355): a randomised, placebo-controlled, double-blind, phase 3 clinical trial. Lancet 396, 1817–1828 (2020).

55 Cortes, J. et al. Pembrolizumab plus chemotherapy in advanced triple-negative breast cancer. N Engl J Med 387, 217–226 (2022).

56 Patwari, A. et al. PD-L2 Landscape and Correlation with Outcome: An Immunomic Analysis. JCO Oncol. Adv. 3 (2026).

57 Behringer, R., Gertsenstein, M., Nagy, K. V. & Nagy, A. Manipulating the mouse embryo: a laboratory manual. (Cold Spring Harbor Laboratory Press, 2014).

58 Hashimoto, M., Yamashita, Y. & Takemoto, T. Electroporation of Cas9 protein/sgRNA into early pronuclear zygotes generates non-mosaic mutants in the mouse. Dev. Biol. 418, 1–9 (2016).

59 Schraml, B. U. et al. Genetic tracing via DNGR-1 expression history defines dendritic cells as a hematopoietic lineage. Cell 154, 843–858 (2013).

60 Kam, J. W. et al. RGMB and neogenin control cell differentiation in the developing olfactory epithelium. Development 143, 1534–1546 (2016).

61 Komuczki, J. et al. Fate-Mapping of GM-CSF Expression Identifies a Discrete Subset of Inflammation-Driving T Helper Cells Regulated by Cytokines IL-23 and IL-1beta. Immunity 50, 1289–1304 e1286 (2019).

62 Lee, R. A., Mao, C., Vo, H., Gao, W. & Zhong, X. Fluorescence tagging and inducible depletion of PD-L2-expressing B-1 B cells in vivo. Ann. N. Y. Acad. Sci. 1362, 77–85 (2015).

63 Hogquist, K. A. et al. T cell receptor antagonist peptides induce positive selection. Cell 76, 17–27 (1994).

64 Stoeckius, M. et al. Cell Hashing with barcoded antibodies enables multiplexing and doublet detection for single cell genomics. Genome Biol. 19, 224 (2018).

65 Rodrigues, P. F. et al. Progenitors of distinct lineages shape the diversity of mature type 2 conventional dendritic cells. Immunity 57, 1567–1585 e1565 (2024).

66 Wolf, F. A., Angerer, P. & Theis, F. J. SCANPY: large-scale single-cell gene expression data analysis. Genome Biol. 19, 15 (2018).

67 Levine, J. H. et al. Data-Driven Phenotypic Dissection of AML Reveals Progenitor-like Cells that Correlate with Prognosis. Cell 162, 184–197 (2015).

68 McInnes, L., Healy, J. & Melville, J. Umap: Uniform manifold approximation and projection for dimension reduction. arXiv:1802.03426 (2018).

69 Finak, G. et al. MAST: a flexible statistical framework for assessing transcriptional changes and characterizing heterogeneity in single-cell RNA sequencing data. Genome Biol. 16, 278 (2015).

70 Moran, A. E. et al. T cell receptor signal strength in Treg and iNKT cell development demonstrated by a novel fluorescent reporter mouse. J. Exp. Med. 208, 1279–1289 (2011).

71 Jennings, E. et al. Nr4a1 and Nr4a3 Reporter Mice Are Differentially Sensitive to T Cell Receptor Signal Strength and Duration. Cell Rep. 33, 108328 (2020).

72 Collins, S. et al. Opposing regulation of T cell function by Egr-1/NAB2 and Egr-2/Egr-3. Eur. J. Immunol. 38, 528–536 (2008).

73 van Dijk, D. et al. Recovering Gene Interactions from Single-Cell Data Using Data Diffusion. Cell 174, 716–729 e727 (2018).

74 Palla, G. et al. Squidpy: a scalable framework for spatial omics analysis. Nature methods 19, 171–178 (2022).

75 Ester, M., Kriegel, H.-P., Sander, J. & Xu, X. in kdd. 226–231.

76 Lun, A. T. L. et al. EmptyDrops: distinguishing cells from empty droplets in droplet-based single-cell RNA sequencing data. Genome Biol 20, 63 (2019).

77 Gayoso, A. & Shor, J. JonathanShor/DoubletDetection: doubletdetection v4. 3.0. post1. Zenodo (2025).

78 Xu, C. et al. Automatic cell-type harmonization and integration across Human Cell Atlas datasets. Cell 186, 5876–5891 e5820 (2023).

79 Dominguez Conde, C., et al. Cross-tissue immune cell analysis reveals tissue-specific features in humans. Science 376, eabl5197 (2022).

