## Supplementary materials for "Dendritic cell PD-L2 restrains intratumoral CD8^+^ T cell immunity"


This file includes:

Extended Data Fig. 1 to 10

Supplementary Fig. 1 to 3

Supplementary Table 1 to 4

**Extended Data Fig. 1**

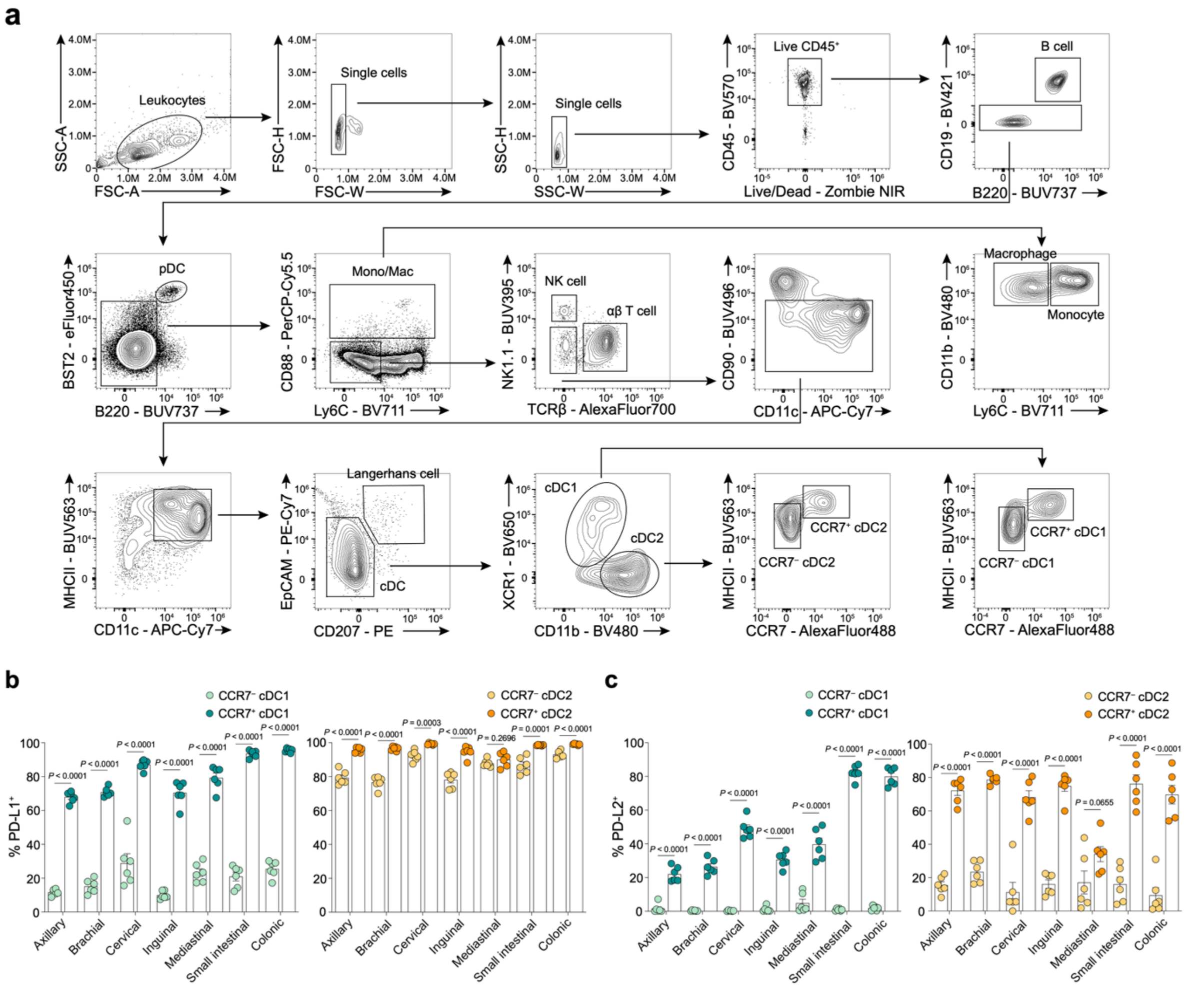


**Extended Data Fig. 1 | High expression of PD-L2 in CCR7+ cDC subsets in steady-state LN. a**, Gating strategy for various immune subsets including CCR7+ cDC1 and CCR7+ cDC2s. **b**, Summary graphs of the frequency of PD-L1+ cells among cDC subsets across LNs from C57Bl/6 mice (n = 6). **c**, Summary graphs of the frequency of PD-L2+ cells among cDC subsets across LNs from C57Bl/6 mice (n = 6). Data in **b**, **c** are representative of two independent experiments. Error bars: mean ± s.e.m.; two-tailed unpaired *t*-test (**b**, **c**). All *P* values are indicated in the corresponding graphs.

**Extended Data Fig. 2**

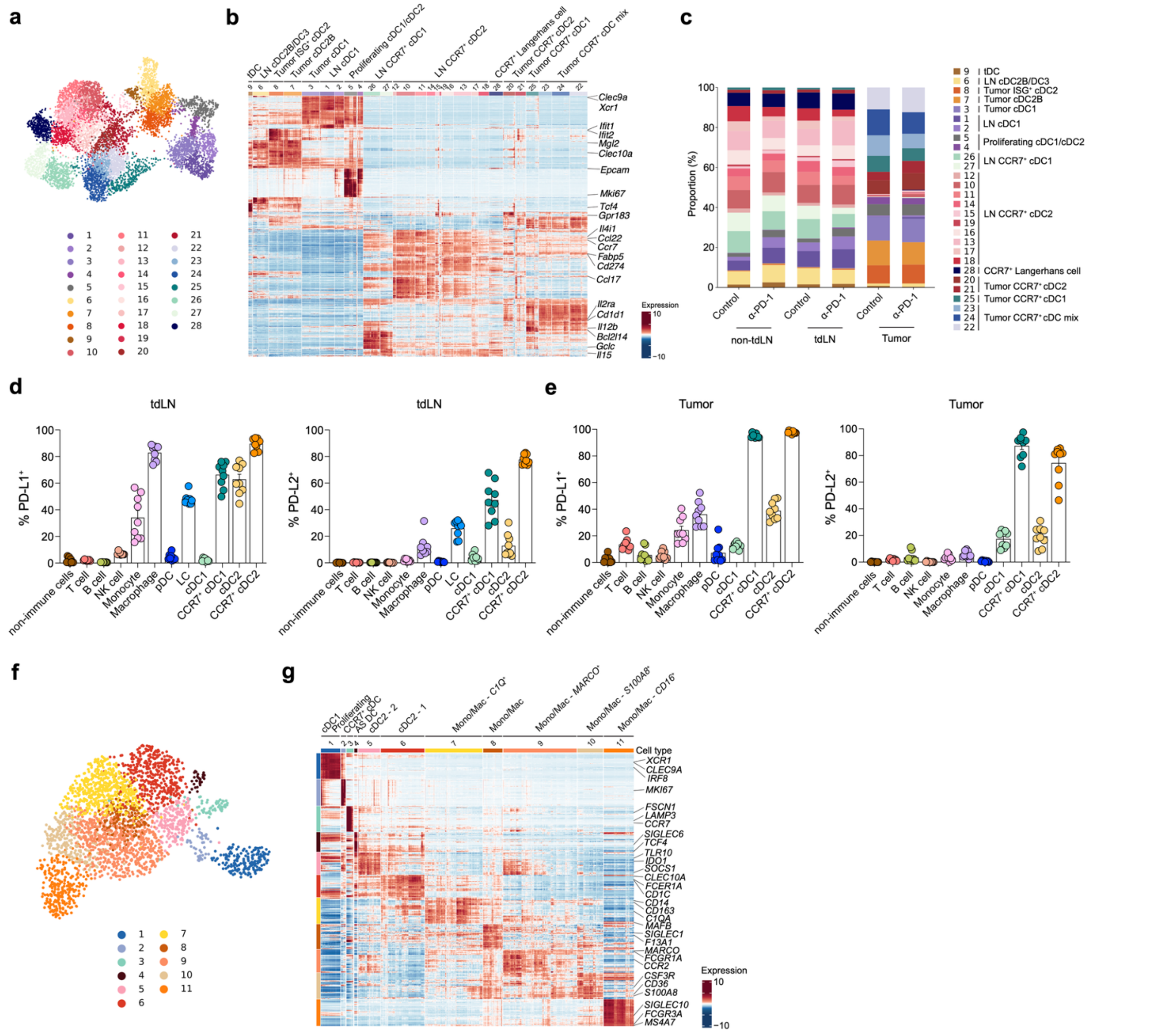


**Extended Data Fig. 2 | Single-cell characterization of dendritic-cell states and PD-1 ligand expression in B16-OVA tumor context. a**–**c**, scRNA-seq analysis of Lin$^{-}$CD11c$^{+}$MHCII$^{+}$ DCs isolated from non-tumor-draining lymph nodes (non-tdLNs), tumor-draining lymph nodes (tdLNs) and tumors of B16-OVA-bearing C57BL/6 mice treated with anti–PD-1 antibody or isotype control (n = 5 mice per group). **a**, UMAP colored by Leiden cluster. **b**, Heatmap showing scaled, imputed expression of the top 50 differentially expressed genes for each cluster (one versus the rest, fold change (FC) > 1.5, adjusted $P < 0.01$). **c**, Distribution of cell types as in (**a**) across non-tdLNs, tdLNs, and tumors from anti–PD-1 or isotype control treated mice. **d**, **e**, C57BL/6 mice were implanted with B16-OVA, and CD45$^{+}$ immune and CD45$^{-}$ non-immune cells were analyzed by flow cytometry (n = 9). Summary graphs of the frequency of PD-L1$^{+}$ and PD-L2$^{+}$ cells among the indicated subsets in (**d**) tdLNs and (**e**) tumors. **f, g**, scRNA-seq on Lin(CD3, CD19, CD56)$^{-}$CD11C$^{+}$HLA-DR$^{+}$ cells isolated from human melanoma. **f**, UMAP colored by Leiden cluster. **g**, Heatmap showing scaled, imputed expression of the top 50 differentially expressed genes for each cluster (one versus the rest, fold change (FC) > 1.5, adjusted $P < 0.01$). Data in **d**, **e** are pooled from two independent experiments.

**Extended Data Fig. 3**

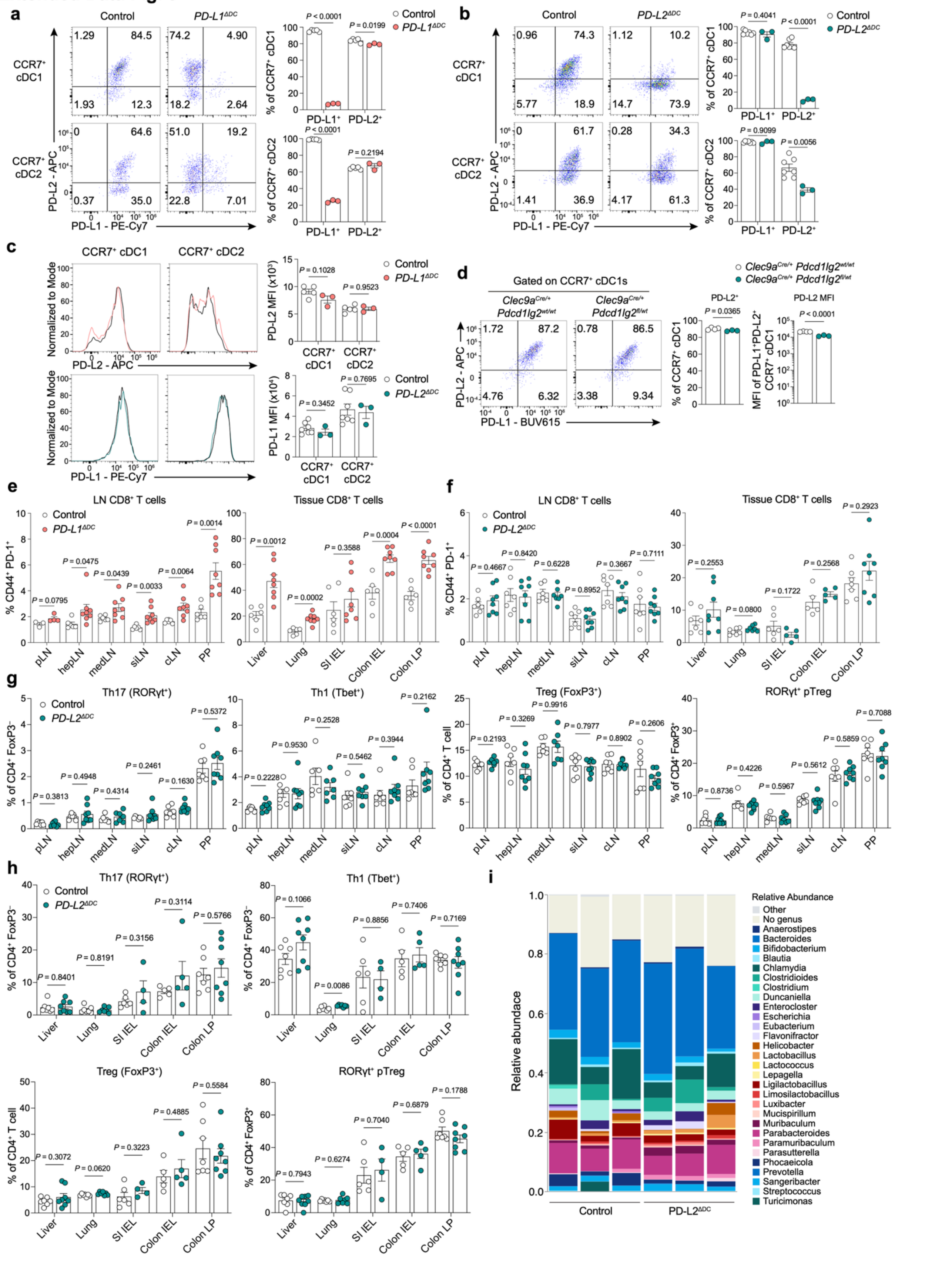

**Extended Data Fig. 3 | Validation of cDC-specific PD-1 ligand deletion and steady-state homeostasis.** **a**, Representative flow plots and summary graphs of the frequency of PD-L1+ and PD-L2+ cells among CCR7+ cDC subsets in colonic LNs from *PD-L1*$^{ΔDC}$ (*n* = 3) and control (n = 5) mice. **b**, Representative flow plots and summary graphs of the frequency of PD-L1+ and PD-L2+ cells among CCR7+ cDC subsets in colonic LNs from *PD-L2*$^{ΔDC}$ (*n* = 3) and control (n = 7) mice. **c**, Representative flow plots and summary graphs of PD-L2 expression by CCR7+ cDC subsets in colonic LNs from *PD-L1*$^{ΔDC}$ (*n* = 3) and control (n = 5) mice (upper), and PD-L1 expression by CCR7+ cDC subsets in colonic LNs from *PD-L2*$^{ΔDC}$ (*n* = 3) and control (n = 7) mice (lower). **d**, Representative flow plots and summary graphs of PD-L2 expression by CCR7+ cDC1s in colonic LNs from *Clec9a*$^{Cre/+}$ *Pdcd1lg2*$^{fl/wt}$ (n = 3) and *Clec9a*$^{Cre/+}$ *Pdcd1lg2*$^{wt/wt}$ (n = 4) mice. **e**, Summary graphs of the frequency of CD44+PD-1+ cells among CD8+ T cells across LNs and tissues from *PD-L1*$^{ΔDC}$ (*n* = 8) and control (n = 6) mice, with exact sample sizes indicated by individual data points; pLN (peripheral LN), hepLN (hepatic LN), medLN (mediastinal LN), siLN (small intestinal LN), cLN (colonic LN), PP (Peyer's Patch), SI IEL (Small intestinal intraepithelial lymphocyte), Colon IEL (Colon intraepithelial lymphocyte), Colon LP (Colon lamina propria). **f**, Summary graphs of the frequency of CD44+PD-1+ cells among CD8+ T cells across LNs and tissues from *PD-L2*$^{ΔDC}$ (*n* = 8) and control (n = 7) and mice, with exact sample sizes indicated by individual data points. **g**, **h**, Summary graphs of the frequency of Th17, Th1, Treg, and RORgt+ pTreg from *PD-L2*$^{ΔDC}$ (*n* = 8) and control (n = 7) mice across LNs (**g**) and tissues (**h**). **i**, Microbiome composition in *PD-L2*$^{ΔDC}$ (n = 3) and control (n = 3) mice. Data in **a**–**d** are representative of two independent experiments. Data in **e–h** are pooled from two independent experiments. Error bars: mean ± s.e.m.; two-tailed unpaired *t*-test (**a**-**h**). All *P* values are indicated in the corresponding graphs.

### Extended Data Fig. 4

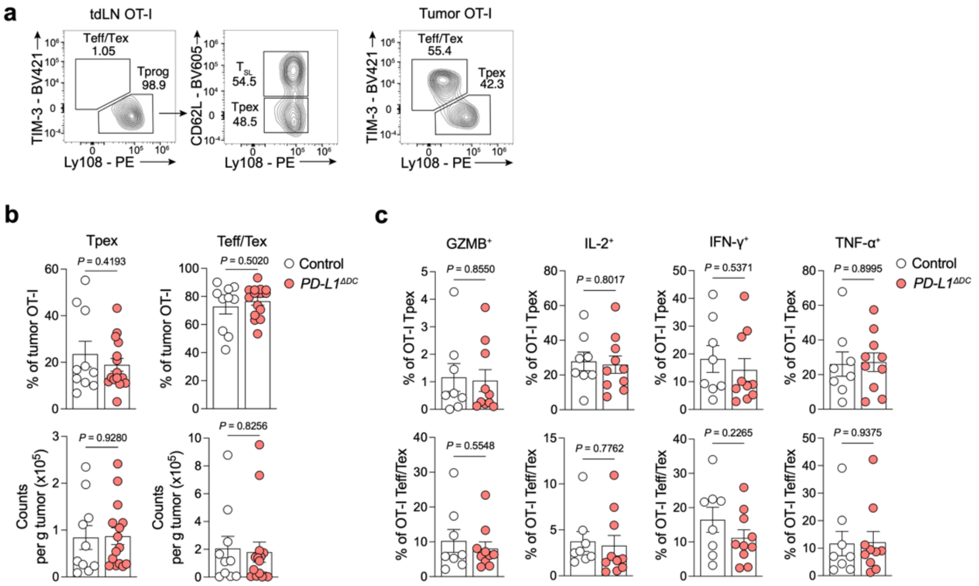


**Extended Data Fig. 4 | Ablation of PD-L1 on cDCs is insufficient to drive effector differentiation within tumors**. **a**, Gating strategy for stem-like ($T_{SL}$), progenitor exhausted (Tpex), and effector/exhausted (Teff/Tex) cells among OT-I cells across tdLNs and tumors. **b**, Summary graphs of the frequency and number of Tpex and Teff/Tex among tumor-infiltrating OT-I cells from *PD-L1$^{ΔDC}$* ($n$ = 15) and control (n = 10) mice. **c**, Summary graphs of the frequency of GZMB+, IL-2+, IFN-γ+ and TNF-α+ cells among tumor-infiltrating OT-I Tpex and Teff/Tex cells from *PD-L1$^{ΔDC}$* (n = 10) and control (n = 8) mice. Data in **b** are pooled from three independent experiments; data in **c** are pooled from two independent experiments. Error bars: mean ± s.e.m.; two-tailed unpaired *t*-test (**b**, **c**). All *P* values are indicated in the corresponding graphs.

### Extended Data Fig. 5

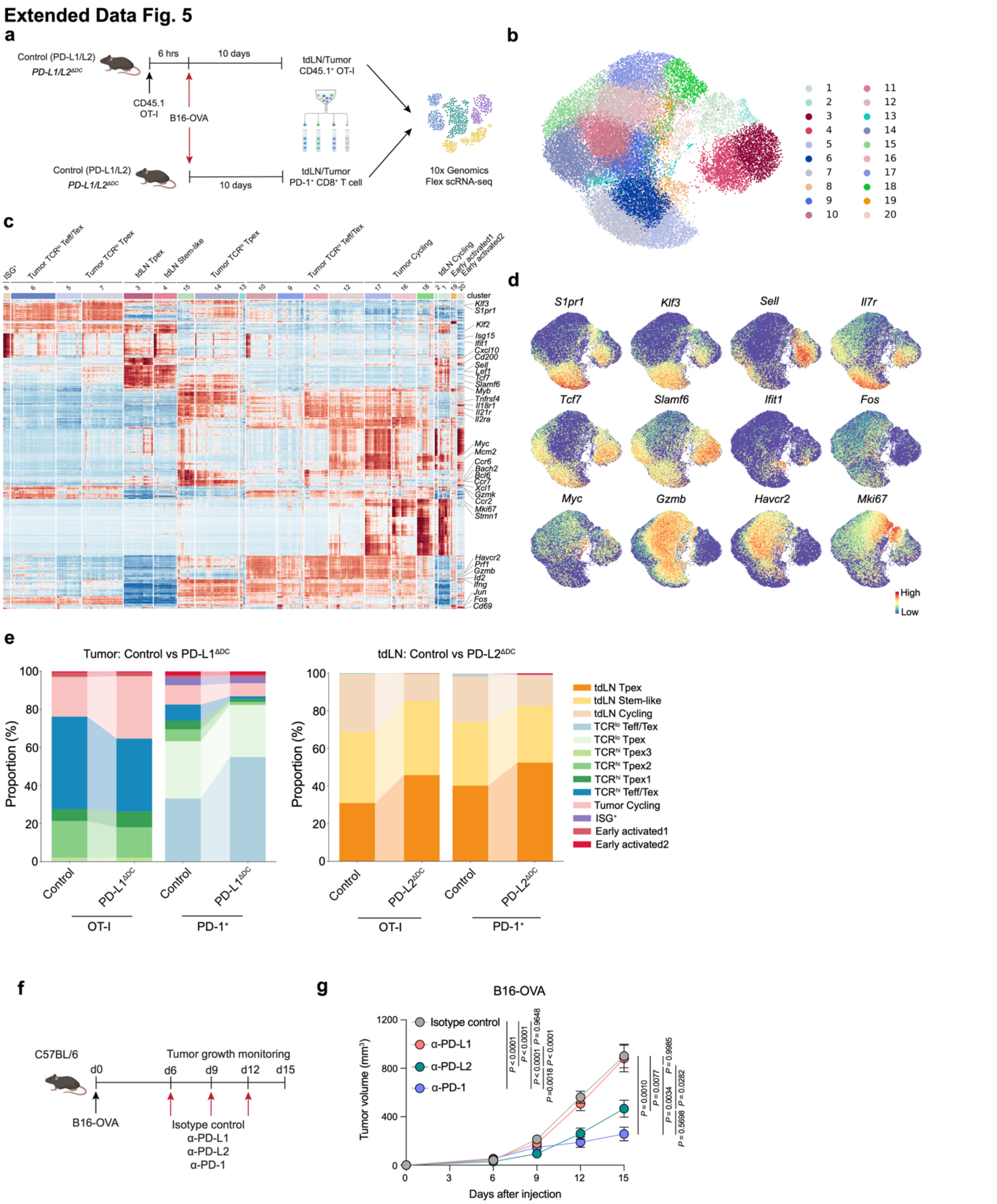

**Extended Data Fig. 5 | Single-cell annotation of CD8+ T cell states and PD-1 ligand perturbation in B16-OVA melanoma**. **a**, Schematic of the experimental design for CD8+ T cell scRNA-seq. **b**, UMAP colored by Leiden cluster. **c**, Heatmap showing scaled, imputed expression of the top 50 differentially expressed genes for each cluster (one versus the rest, fold change (FC) > 1.5, adjusted *P* < 0.01). **d**, UMAP colored by unimputed expression of indicated marker genes. **e**, Distribution of CD8+ T cell types in tumors from PD-L1 control and *PD-L1$^{ΔDC}$* mice (left) or tdLNs from PD-L2 control and *PD-L2$^{ΔDC}$* mice (right). **f**, Schematic of the experimental design. **g**, Tumor growth curves in mice implanted with B16-OVA and treated with anti-PD-L1 (n = 8), anti-PD-L2 (n = 9), anti-PD-1 (n = 5) or isotype control (n = 13). Panel **a** and **f** were created using BioRender; https://biorender.com. Data in **g** are pooled from five independent experiments. Error bars: mean ± s.e.m.; one-way ANOVA and two-way ANOVA (**g**). All *P* values are indicated in the corresponding graphs.

### Extended Data Fig. 6

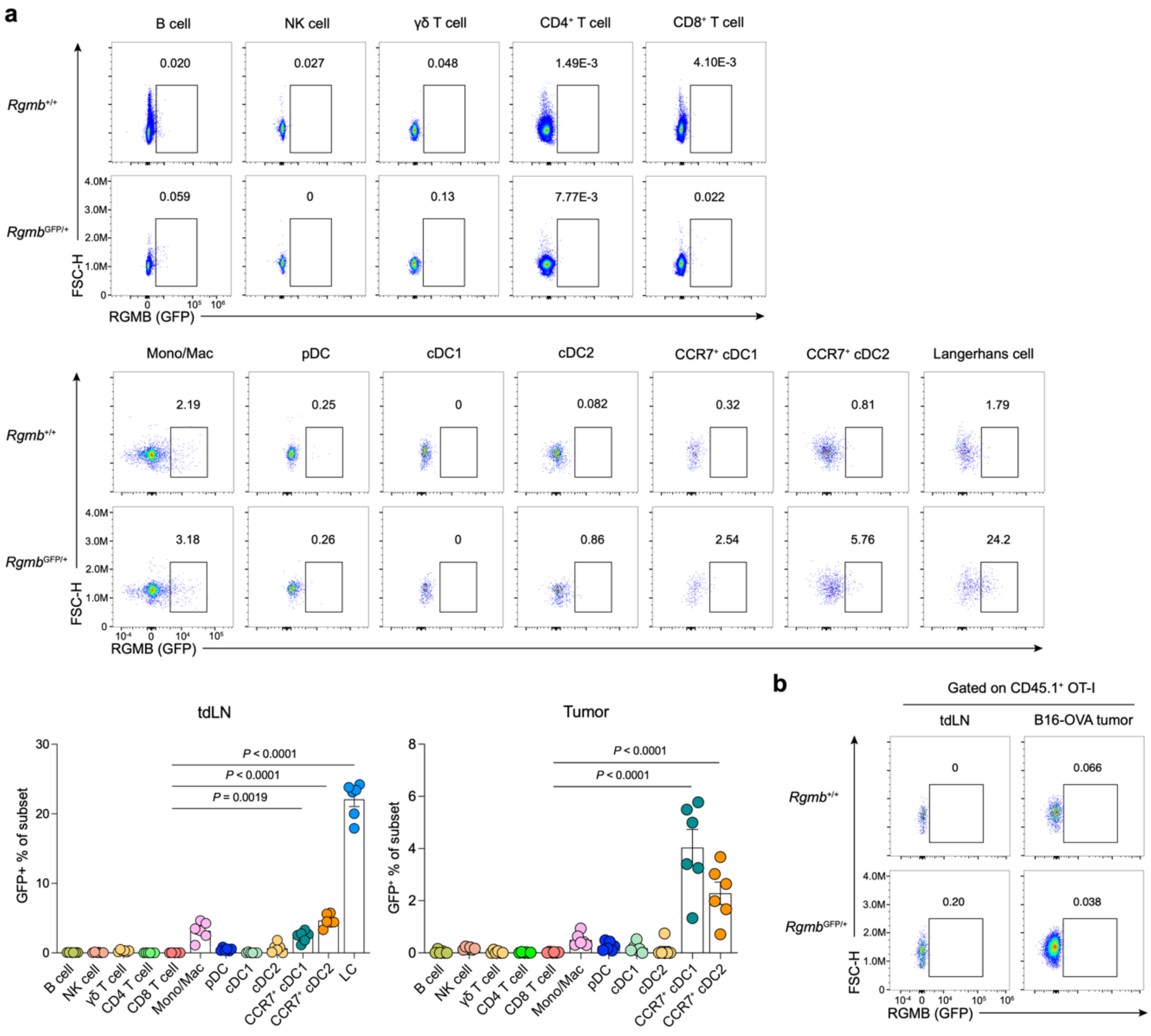


**Extended Data Fig. 6 | RGMb expression in immune and antigen-specific CD8+ T cells in B16-OVA-bearing mice.** **a**, Representative flow plots and summary graphs of the frequency of RGMb(GFP)+ cells among the indicated cell types in tdLNs and tumors of B16-OVA-bearing *Rgmb*$^{GFP/+}$ (n = 6) or control *Rgmb*$^{+/+}$ mice. **b**, Congenically marked *Rgmb*$^{GFP/+}$ or *Rgmb*$^{+/+}$ OT-I cells were adoptively transferred into C57Bl/6 mice 6 hours prior to B16-OVA implantation and analyzed in tdLNs and tumors 10 days later by flow cytometry. Representative flow plots of the frequency of RGMb+ cells among OT-I cells in tdLNs and tumors. Data in **a** are pooled from two independent experiments. Data in **b** are representative of two independent experiments. Error bars: mean ± s.e.m.; one-way ANOVA (**a**). All *P* values are indicated in the corresponding graphs.

**Extended Data Fig. 7**

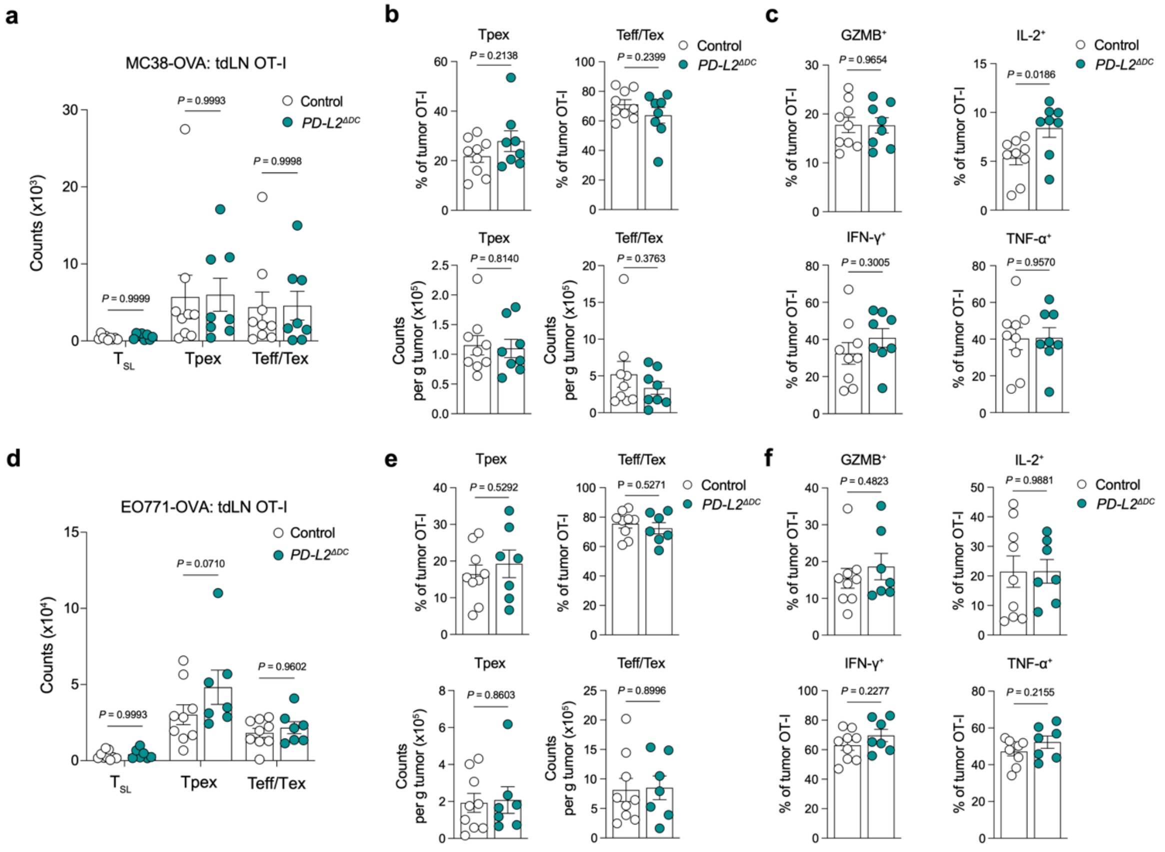


**Extended Data Fig. 7 | Limited effects of cDC-specific PD-L2 ablation on tumor-specific CD8⁺ T cell responses in MC38-OVA and EO771-OVA tumors. a-f**, Congenically marked OT-I cells were adoptively transferred into *PD-L2$^{ΔDC}$* or control mice 6 hours prior to MC38-OVA or EO771-OVA implantation and analyzed in tdLNs and tumors 10 days later by flow cytometry. **a**, Number of OT-I $T_{SL}$, Tpex, and Teff/Tex cells in tdLNs from MC38-OVA-bearing *PD-L2$^{ΔDC}$* (n = 8) or control (n = 9) mice. **b**, Summary graphs of the frequency and number of OT-I Tpex and Teff/Tex cells in tumors from MC38-OVA-bearing *PD-L2$^{ΔDC}$* (n = 8) or control (n = 9) mice. **c**, Summary graphs of the frequency of GZMB$^+$, IL-2$^+$, IFN-γ$^+$ and TNF-α$^+$ cells among tumor-infiltrating OT-I cells from MC38-OVA-bearing *PD-L2$^{ΔDC}$* (n = 8) or control (n = 9) mice. **d**, Number of OT-I $T_{SL}$, Tpex, and Teff/Tex cells in tdLNs from EO771-OVA-bearing *PD-L2$^{ΔDC}$* (n = 7) or control (n = 9) mice. **e**, Summary graphs of the frequency and number of OT-I Tpex and Teff/Tex cells in tumors from EO771-OVA-bearing *PD-L2$^{ΔDC}$* (n = 7) or control (n = 9) mice. **f**, Summary graphs of the frequency of GZMB$^+$, IL-2$^+$, IFN-γ$^+$ and TNF-α$^+$ cells among tumor-infiltrating OT-I cells from EO771-OVA-bearing *PD-L2$^{ΔDC}$* (n = 7) or control (n = 9) mice. Data in **a–f** are pooled from two independent experiments. Error bars: mean ± s.e.m.; two-way ANOVA (**a**, **d**), two-tailed unpaired *t*-test (**b**, **c**, **e**, **f**). All *P* values are indicated in the corresponding graphs.

### Extended Data Fig. 8

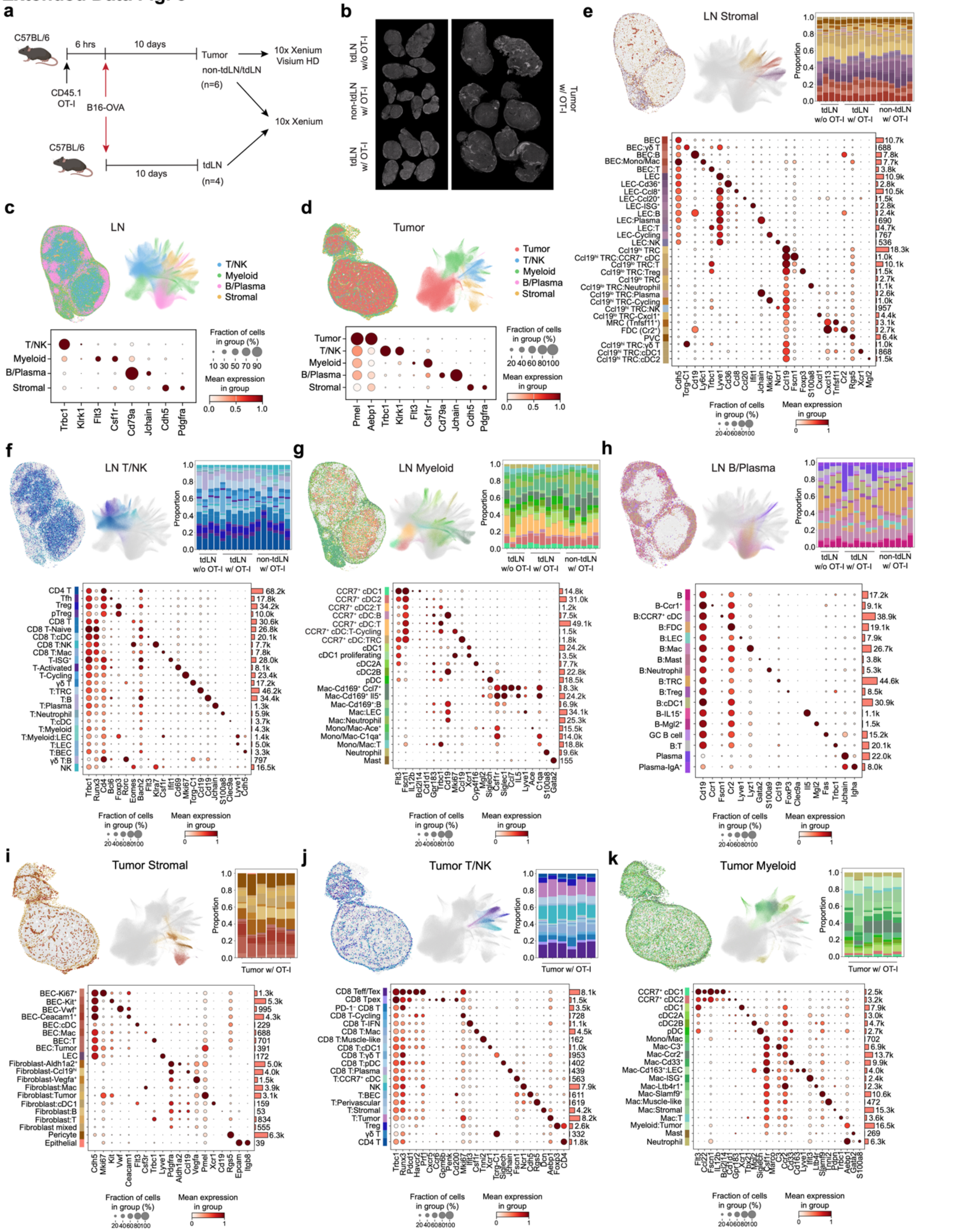

**Extended Data Fig. 8 | 10x Xenium spatial transcriptomics on LNs and tumors of B16-OVA-bearing mice.** **a**, Schematic of the spatial transcriptomics experimental design. **b**, Spatial orientation of LN and tumor samples. **c**, **d**, Representative spatial plots and UMAPs of broad cell lineages in LN (**c**) and tumor (**d**) samples, with dot plots showing lineage-specific marker expression. **e–h**, Representative spatial plots, UMAPs, cell-type composition for each individual sample and dot plots showing subset-specific marker expression in LN stromal subsets (**e**), T/NK subsets (**f**), myeloid subsets (**g**) and B/plasma subsets (**h**). **i–k**, Representative spatial plots, UMAPs, cell-type composition for each individual sample and dot plots showing subset-specific marker expression in tumor stromal subsets (**i**), T/NK subsets (**j**) and myeloid subsets (**k**). Panel **a** was created using BioRender; https://biorender.com. UMAPs and dot plots in **c–k** were generated from pooled LNs or tumors, as indicated in **b**. Inherent constraints of cell boundary segmentation resulted in transcript mixing from proximal cells, yielding mixed cell type annotations, indicated by a colon in panels **e–k**, e.g. cells annotated as "CD8 T:cDC1" expressed markers of both CD8 T cells and cDC1s.

### Extended Data Fig. 9

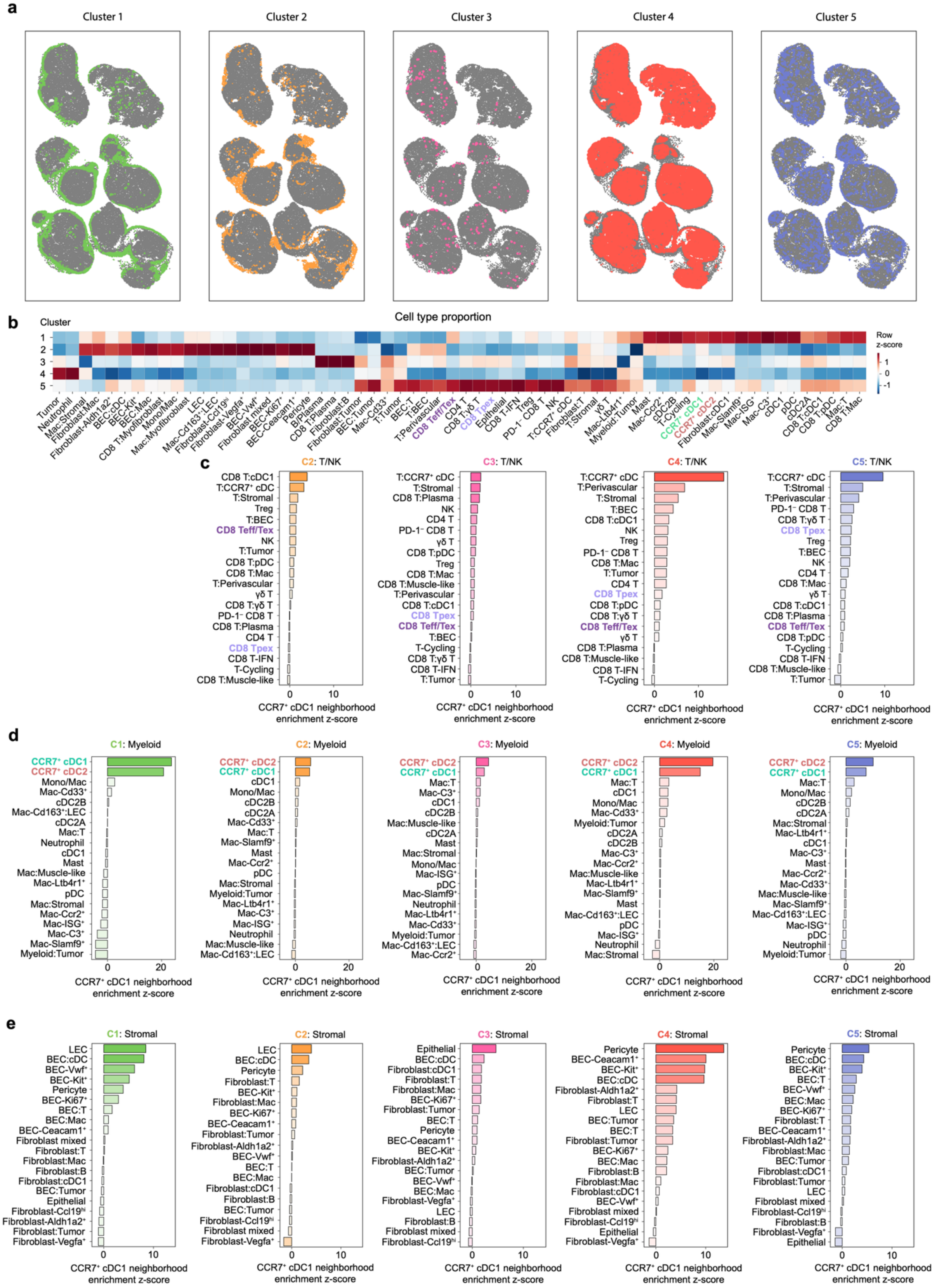

**Extended Data Fig. 9 | Spatial organization of CCR7+ cDC1-associated neighborhoods in B16-OVA tumors. a**, Spatial representation of tumor neighborhood clusters. **b**, Heatmap showing cell subtype enrichment across neighborhood clusters. **c–e**, Summary graphs showing enrichment z-scores of tumor T/NK (**c**), myeloid (**d**) and stromal (**e**) subsets around CCR7+ cDC1 neighborhoods across clusters. Data in **b–e** were generated from pooled tumors.

### Extended Data Fig. 10

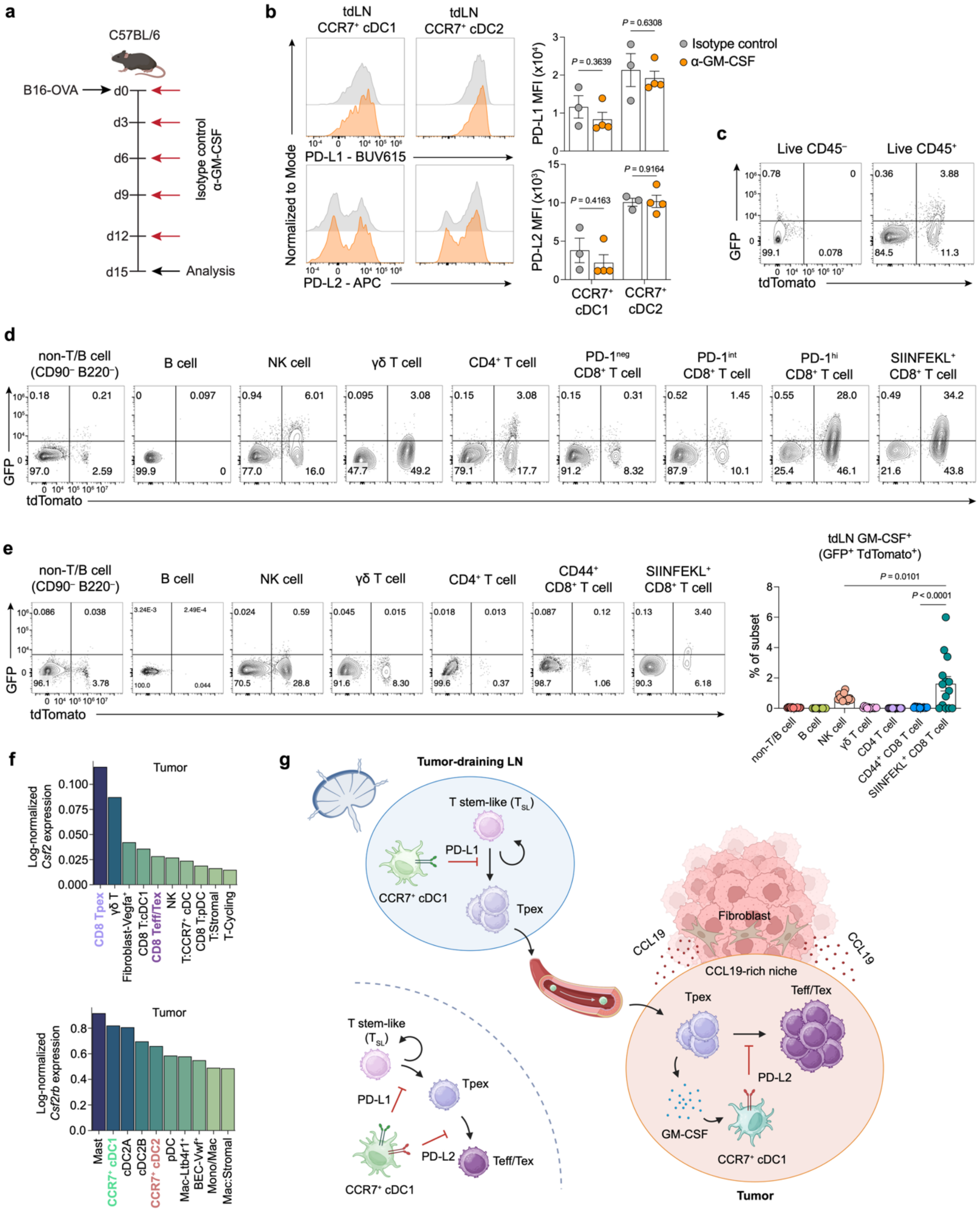

**Extended Data Fig. 10 | Cellular mapping of GM-CSF sources and receptor expression in B16-OVA tumors. a**, Schematic of the GM-CSF blockade experimental design. **b**, Representative flow plots and summary graphs of PD-L1 and PD-L2 expression by CCR7+ cDC subsets in tdLNs from anti–GM-CSF (n = 4) or isotype control (n = 3) treated mice. **c-e**, $Csf2^{icre\text{-}EGFP}$ $R26^{\mathrm{lsl\text{-}tdTomato}}$ ($\mathrm{FROG}^{\mathrm{Ai14}}$) mice were implanted with B16-OVA and analyzed at day 14. **c**, Representative flow plots of the frequency of GM-CSF+ (GFP+tdTomato+) cells among CD45− non-immune and CD45+ immune cells in tumors. **d**, Representative flow plots of the frequency of GM-CSF+ (GFP+tdTomato+) cells among the indicated cell types in tumors. **e**, Representative flow plots and summary graphs of the frequency of GM-CSF+ (GFP+tdTomato+) cells among the indicated cell types in tdLNs (n = 14 mice). **f**, Summary graphs of the expression of *Csf2* and *Csf2rb* among cell types in the tumors defined from 10x Xenium data. **g**, Schematic delineating the functionally and spatially distinct roles of DC PD-L2 in shaping CD8+ T cell differentiation within intratumoral niches. Panel **a** and **g** were created using BioRender; https://biorender.com. Data in **b** are representative of two independent experiments. Data in **c**, **d** are representative of three independent experiments. Data in **e** are pooled from three independent experiments. Error bars: mean ± s.e.m.; two-tailed unpaired *t*-test (**b**), one-way ANOVA (**e**). All *P* values are indicated in the corresponding graphs.

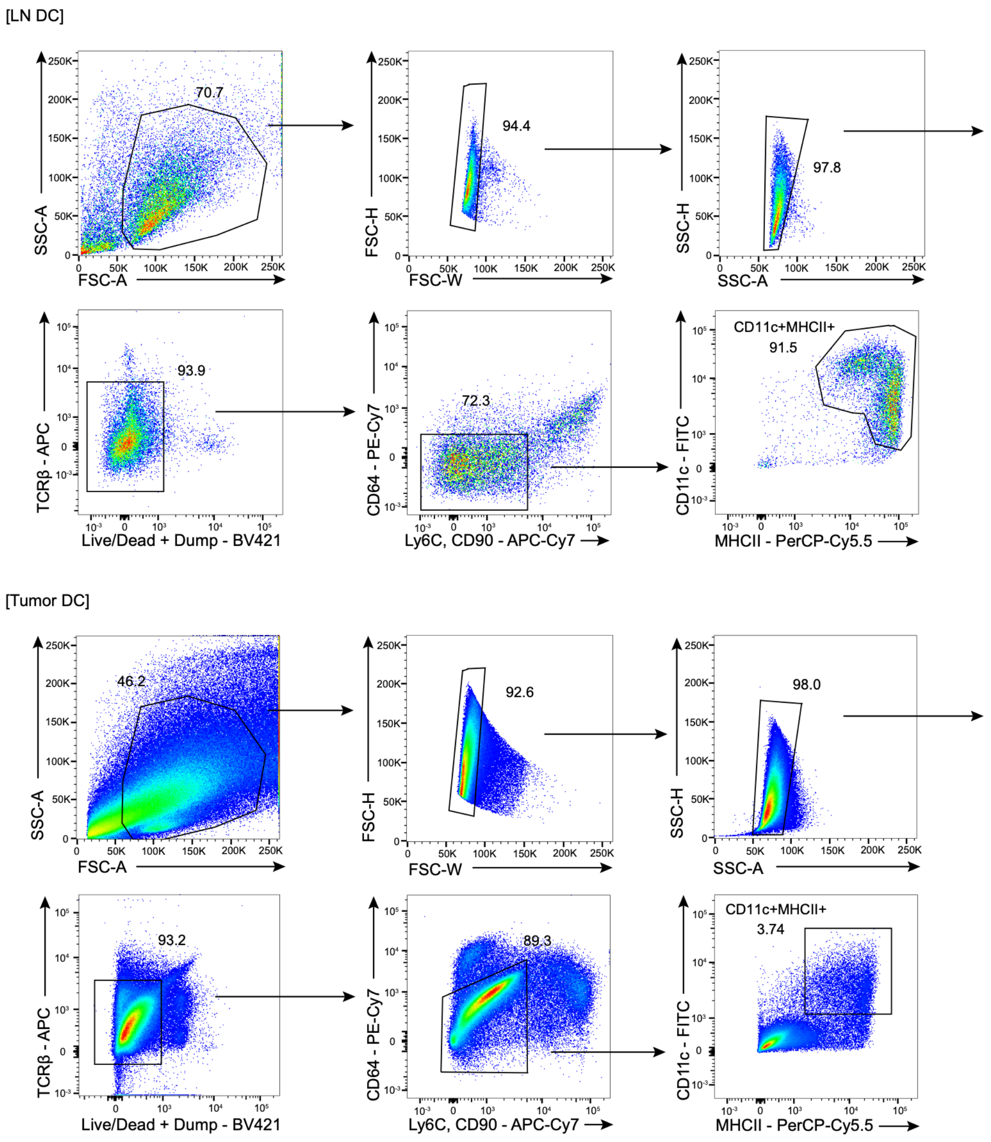


**Supplementary Fig. 1 | FACS isolation of DCs from LNs and tumors of B16-OVA-bearing mice.** Representative gating strategy for the isolation of Lin$^-$(CD3$^-$CD19$^-$CD90$^-$)CD64$^-$Ly6C$^-$CD11c$^+$MHCII$^+$ DCs from LNs and tumors of B16-OVA-bearing mice. Dump: NK1.1, TCRγδ.

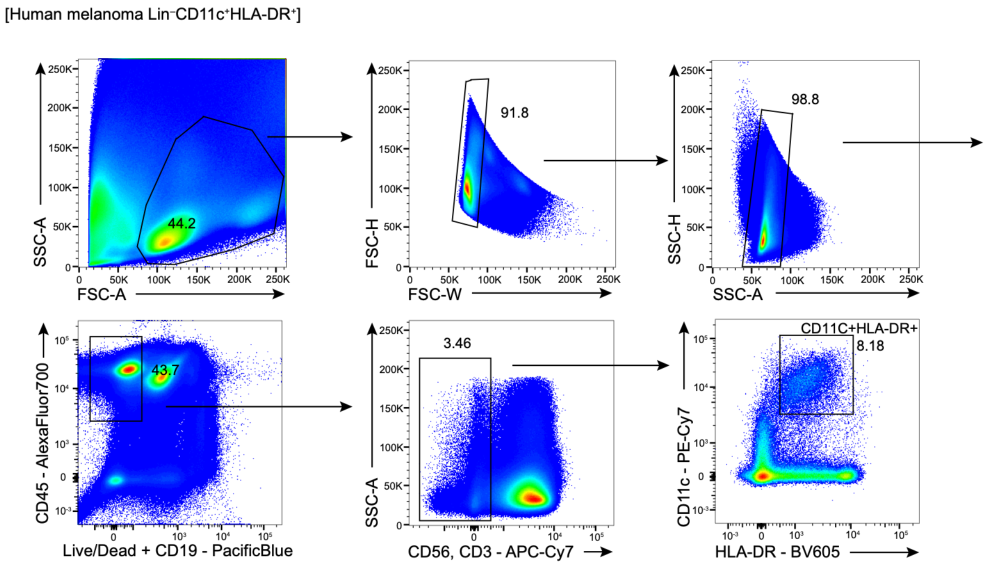


**Supplementary Fig. 2 | FACS isolation of Lin⁻CD11c⁺HLA-DR⁺ cells from human melanoma.** Representative gating strategy for the isolation of Lin⁻(CD3⁻CD19⁻CD56⁻)CD11c⁺HLA-DR⁺ cells from human melanoma.

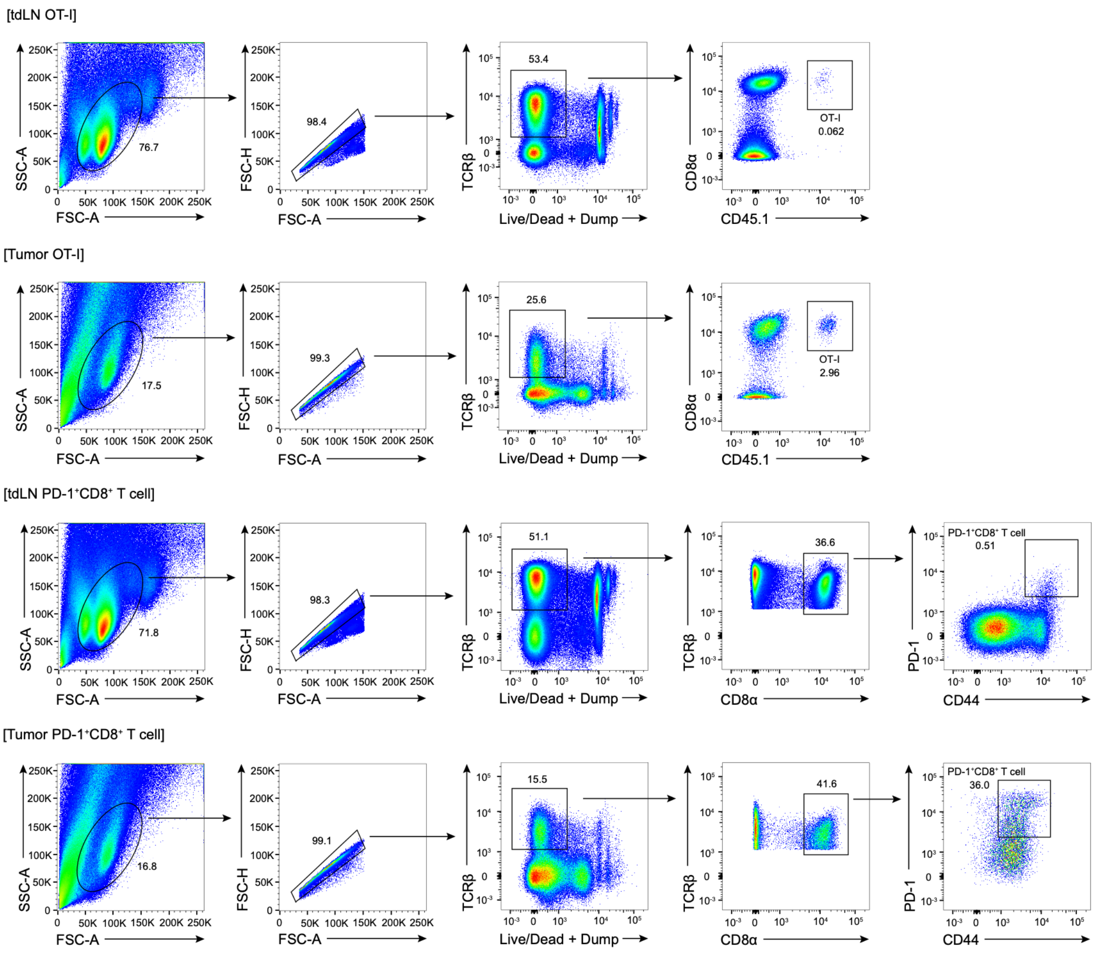


**Supplementary Fig. 3 | FACS isolation of OT-I cells and endogenous PD-1$^+$CD8$^+$ T cells from tdLNs and tumors of B16-OVA-bearing mice.** Representative gating strategy for the isolation of congenic OT-I cells (TCRβ$^+$CD8α$^+$CD45.1$^+$) and endogenous PD-1$^+$CD8$^+$ T cells (TCRβ$^+$CD8α$^+$CD44$^+$PD-1$^+$) from tdLNs and tumors of B16-OVA-bearing mice. Dump: NK1.1, TCRγδ, CD1d tetramer.

**Supplementary Table 1 |** A customized 472-gene panel, designed to identify immune, stromal and tumor cell populations.

**Supplementary Table 2 |** CCR7$^{+}$ cDC1/Tpex-associated stromal niche signature genes identified by 10x Visium HD.

**Supplementary Table 3 |** Human melanoma patient clinical information.

**Supplementary Table 4 |** Antibodies used for flow cytometry and FACS.
